# Aging in Fast-Forward: An Inducible SIRT6 Deficiency model as a Lens on Brain Aging and Neurodegeneration

**DOI:** 10.64898/2026.02.15.705977

**Authors:** Yuval Rabuah Botton, Dmitrii Smirnov, Songchen Yang, Daniel Stein, Zeev Slobodnik, Ekaterina Eremenko, Shai Kaluski-Kopatch, Monica Einav, Ekaterina Khrameeva, Debra Toiber

## Abstract

Aging and neurodegeneration occur gradually, making in vitro modeling challenging and costly. We generated a time-resolved, reversible neuron-like aging model by gradually depleting SIRT6. Within three weeks, their transcriptomes recapitulated brain-aging signatures and hallmarks. RNA-seq revealed clusters of nonlinear changes and predicted oscillatory DNA-damage and apoptotic programs, allowing stress response and adaptation. SIRT6 loss led to nuclear envelope breakdown and micronuclei accumulation, resembling accelerated-aging phenotypes. Some defects could be rescued by re-expressing SIRT6. As a tool for discovery, we uncovered disrupted nucleocytoplasmic transport as an aging pathway shared with neurodegenerative disorders. When SIRT6 depletion reached 30 days, transcriptional changes correlated with those in Alzheimer’s patients, but reversed after SIRT6 re-expression. Pathways associated with shSIRT6 and healthy aging became anticorrelated with AD, pointing to critical signatures. Our affordable and easy-to-use model captures key molecular features of aging, distinguishes physiological from disease-linked changes, and accelerates mechanistic discovery in a controllable, neuron-based system.

**Graphical Abstract:**
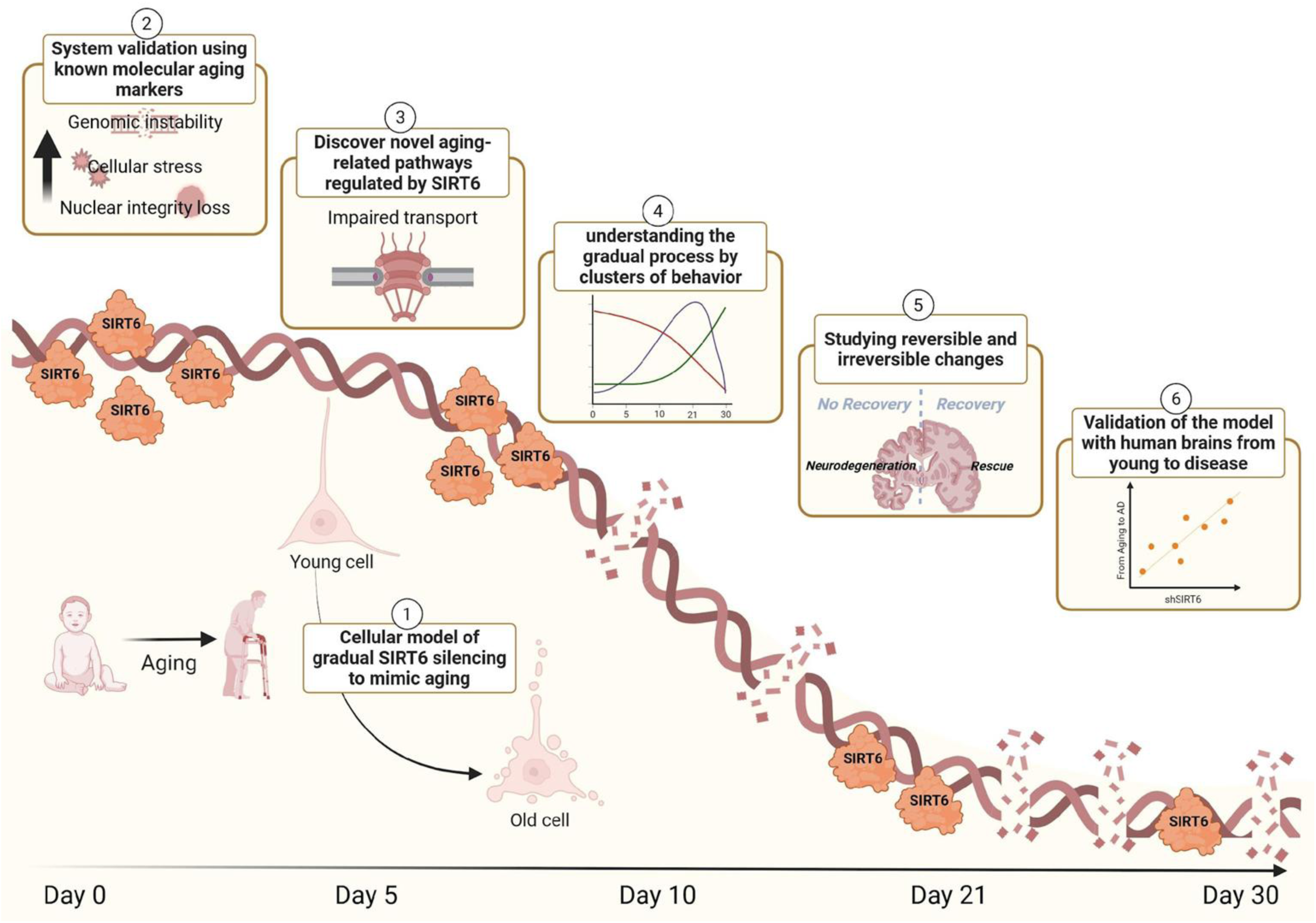
A Novel Reversible Cellular Model of Gradual SIRT6 Decline Reveals Transcriptomic Transitions from Aging to Neurodegeneration. (1) We developed a neuron-like cellular system with gradual, inducible SIRT6 depletion to mimic the aging process. The model enables: (2) Recapitulation of hallmark features of aging, including genomic instability, cellular stress, and nuclear integrity loss. (3) Identification of novel SIRT6-regulated pathways, including impaired nucleocytoplasmic transport. (4) Clustering of dynamic, non-linear gene expression trajectories. (5) Distinction between reversible and irreversible changes following SIRT6 re-expression. (6) Transcriptomic comparison with human brain aging and AD revealing divergence points between healthy and pathological aging.

## Introduction

Aging is characterized by a progressive decline in physiological function, leading to increased susceptibility to diseases such as neurodegeneration^1^. As the world’s population ages, projections indicate that by 2050, the number of people over 60 will nearly double, emphasizing the urgent need to understand the molecular mechanisms underpinning aging (World Health Organization, 2024). Current research has categorized nine molecular and cellular hallmarks of aging, including genomic instability, telomere attrition, and cellular senescence. However, studying these processes in depth requires experimental models that are often costly and time-consuming. Traditional models struggle to capture the complexity of the human aging process, particularly in cellular systems. Many commonly used models compare senescence to aging, or rely on cells deficient in a key protein, missing the graduality and nonlinear changes that occur in the process, especially in a complex organ like the brain.

SIRT6, a member of the sirtuin family of NAD+-dependent deacetylases, has emerged as a key regulator of aging and neurodegeneration^2,3^. Sirtuins are known to influence a variety of cellular processes, including DNA repair, glucose metabolism, and mitochondrial function, all of which are integral to maintaining cellular homeostasis as organisms age^4–8^. Among the sirtuins, SIRT6 stands out due to its role in protecting against genomic instability, regulating gene expression, and maintaining metabolic balance^9–11^. It is primarily involved in histone deacetylation, specifically targeting histone H3 at lysine residues K9, K56, and K18^9^. This activity positions SIRT6 as a critical factor in safeguarding genomic integrity throughout life.

Studies of SIRT6-deficient models provide compelling insights into its function. Mice lacking SIRT6 exhibit premature aging phenotypes, genomic instability, and shortened lifespans, typically dying within weeks of birth^12^. On the other hand, mice overexpressing SIRT6 are long-lived^13^. Importantly, SIRT6-deficient mice exhibit tissue-specific DNA damage, particularly in the brain^14^. As SIRT6 expression decreases with age, its absence in the brain is especially deleterious, exacerbating the onset of neurodegenerative diseases like AD^15–17^. Brain-specific SIRT6 knockout models (brS6KO) demonstrate severe neurodegenerative-like phenotype, such as increased DNA damage, neuronal loss, and the accumulation of hyperphosphorylated and acetylated tau, hallmarks of Alzheimer’s disease (AD) and other neurodegenerative conditions^18,19^. Our previous studies have shown that SIRT6 plays a multifaceted role in maintaining brain health by regulating tryptophan metabolism, transcriptional control, and proteostasis – processes that when disrupted, contribute to neurodegeneration^20–22^. Moreover, we and others have previously shown that SIRT6 levels are significantly reduced in the brains of AD patients, compared to age-matched controls^18,23,24^. Together, this suggests that a lack of SIRT6 in the brain may be especially detrimental, leading to accelerated aging and neurodegeneration, and further reinforcing their connection.

In this work, we aimed to bridge the gap in aging research by developing a neuronal-like model using the SHSY-5Y cell line, where SIRT6 expression is gradually silenced in a doxycycline-inducible manner. This system enabled us to track the dynamic changes in cellular pathways that occur during the aging process. Our model provides a platform to uncover not only novel pathways involved in aging but also the temporal dynamics of their emergence. This approach is crucial for understanding when specific processes are activated during aging, and which may be reversed. Importantly, our model offers a powerful tool for identifying new therapeutic targets that can mitigate or reverse aging-related damage, a critical step toward addressing neurodegenerative diseases.

One of the most striking findings of our study is the strong correlation between the transcriptomic profiles of our gradual SIRT6 silencing model and those of aging and AD. First, we confirmed that our models align with the aging brain in key pathways and hallmarks of aging. Next, we uncovered clusters of transcriptional behavior that exhibit oscillatory patterns, shifting our perspective from progressive changes to a more complex transition. Finally, we found that many of the pathways dysregulated in our model overlapped with those identified in human AD brains, and this correlation increases with our “cellular aging”, further supporting the relevance of SIRT6 in neurodegeneration. Moreover, we demonstrated that re-expressing SIRT6 for a short period (9 days) in our model was sufficient to reverse several key aging-related pathways and significantly reduce the correlation with aged and AD brains. This discovery opens the door to potential therapeutic interventions targeting SIRT6 and the affected pathways, offering hope that some aging processes may be reversible.

## Results

### Generating the Cellular Model for Investigating SIRT6-Driven Aging Dynamics

While most current cellular models for studying aging have focused on senescent cells, these models fail to fully capture the gradual and complex nature of the aging process, particularly in a complex organ like the brain. To address the need for a non-senescent, gradual cellular model system to study aging, we developed a widely used neuronal-like cell line, SH-SY5Y, with inducible expression of shSIRT6 controlled by doxycycline (Dox). The cells were a pooled population, which prevents single-clone effects and allows us to focus on the broad impact of shSIRT6 (Figure 1A). Because SIRT6 is important in aging models, with SIRT6 deletion accelerating aging and high SIRT6 expression slowing it, we chose to gradually reduce SIRT6 levels over a three-week period (Figure 1B). This approach balances sufficiently slow progression with observable gradual changes. A stable inducible cell line was treated for three weeks with increasing doses of Dox to achieve a gradual silencing effect. Following this period, we stopped Dox treatment, allowed the cells to recover SIRT6 levels for nine days, and assessed reversible versus irreversible changes. Various Dox concentrations (0.2 μg/ml, 0.5 μg/ml, 1 μg/ml) were applied weekly, leading to a progressive decline in both SIRT6 mRNA and protein levels, as shown by Real-Time qPCR (Figure 1C), western blot (Figure 1D), and immunofluorescence (Figure 1E, 1F), confirming the gradual reduction in SIRT6 expression.

**Figure 1:**
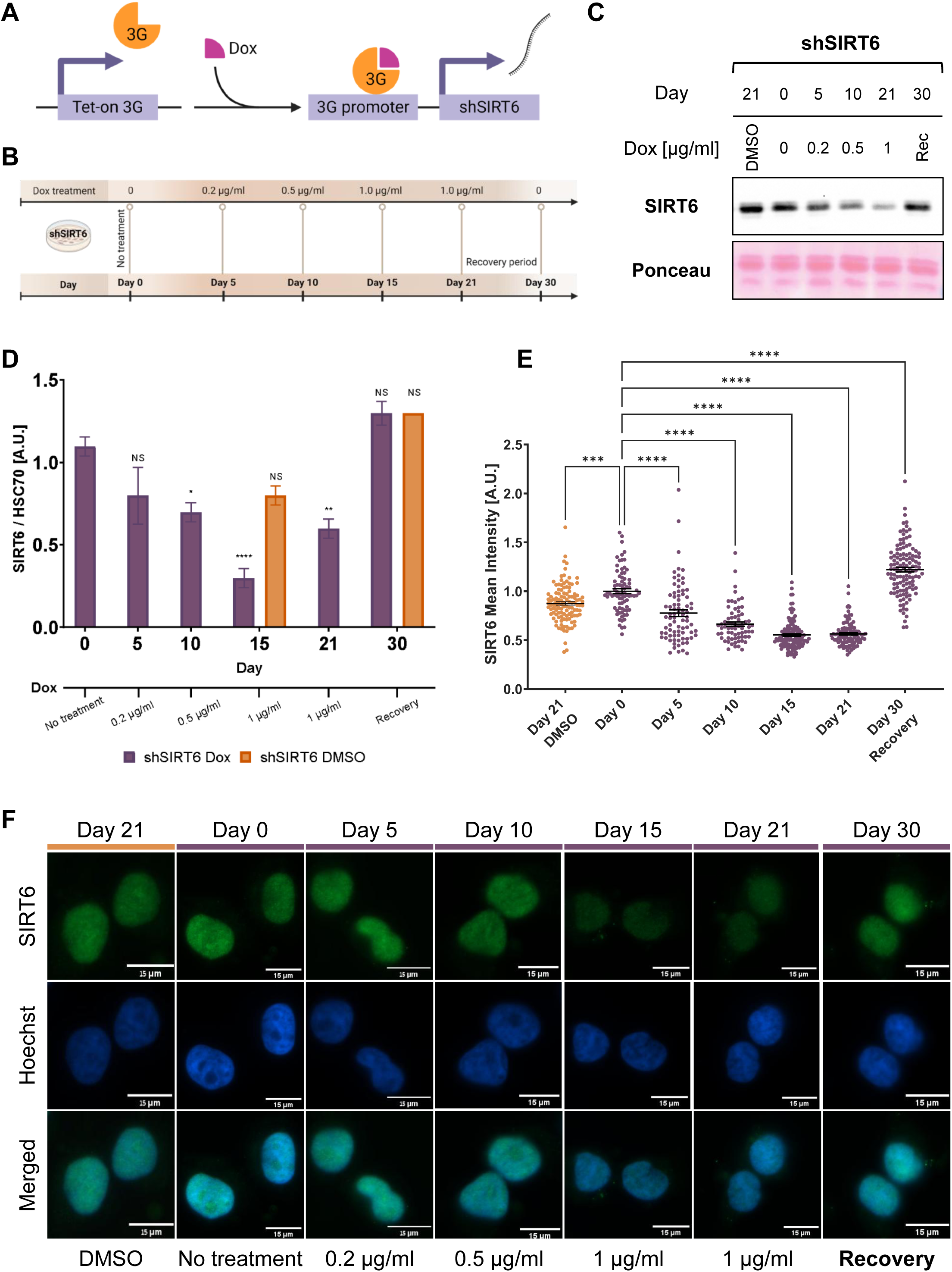
Generating the Cellular Model for Investigating SIRT6-Driven Aging Dynamics. **(A)** The Tet-on transactivator protein is expressed and binds to doxycycline (Dox) to modulate shSIRT6 expression. **(B)** Schematic representation of the aging-mimic model with increasing Dox concentrations, followed by a recovery phase. **(C)** Western blot of SIRT6 protein expression across different Dox concentrations, illustrating the corresponding changes in protein levels. **(D)** Real-time qPCR showing a decrease in SIRT6 mRNA expression with increasing Dox concentrations. **(E)** Quantification of nuclear SIRT6 mean intensity by immunofluorescence, showing a decrease in protein levels across the time course. **(F)** Representative immunofluorescence images corresponding to (E). The Data represents mean ± SEM (ANOVA, *P<0.05, **P<0.01, ***P<0.001, ****P<0.0001).

To gain a comprehensive view of transcriptomic changes in a cellular model, we performed transcriptomic analysis using RNA sequencing (RNA-seq) on cells sampled at distinct time points during Dox treatment (Suppl. Figure 1A). First, considering the potential effects of Dox on gene expression, reported on ^25^, we conducted a comparative analysis between untreated and Dox-treated control cells (shCTRL) (Suppl. Figure 1A). Differentially expressed genes (DEGs) identified from this comparison were attributed to the “Off-target / Dox effect” and excluded from further analysis in our next experiments (Suppl. Figure 1B). Next, we sampled shSIRT6 cells at distinct time points during Dox treatment. To test the side effect of DMSO, we used shSIRT6 with DMSO (mock treatment) cells on day 15, choosing this time point as an intermediate point in the experiment (Suppl. Figure 1C). Principal component analysis (PCA) demonstrated the similarity among replicates, confirming the robustness of our experimental design (Suppl. Figure 1D). As we expected, the number of DEGs between shSIRT6 treated with Dox for 21 days and shCTRL treated with Dox for 15 days was significantly higher than the number of “Dox effect” genes, indicating that SIRT6 silencing drives substantial gene expression changes beyond those induced by Dox alone (Suppl. Figure 1E). To analyze the global changes in gene expression driven by the gradual decline of SIRT6, we performed Pearson correlation analysis of transcriptomic profiles (Suppl. Figure 1F). Hierarchical clustering of samples revealed high consistency among biological replicates, with significant time-dependent gene expression changes primarily driven by the progressive reduction of SIRT6, highlighting its crucial role in regulating gene expression. These results suggest that our transcriptomic data provide a valuable resource for studying the gradual changes associated with SIRT6 decay.

### Dynamic Changes in Aging-Related Pathways in Our Gradual SIRT6 Decline Model Parallel Brain Gradual Changes

To test the relevance of our cellular system as an aging model, we analyzed aging-related pathways, specifically focusing on those recognized as hallmarks of aging (Figure 2A). The tested pathways show the expected expression patterns, for example, an increase in glucose metabolic process (ANOVA p-value = 0.15) and a reduction in mitochondrial function (ANOVA p-value ≤ 0.0001). Re-expression of SIRT6 at Day 30 recovery did not alter the overall trend of certain biological processes, including RNA splicing, ribosome biogenesis, and protein folding, which are commonly affected in aging tissues. The gradual downregulation of these critical pathways, coupled with no elevation trend upon recovery, suggests that they were modulated in a SIRT6-dependent manner and ultimately reached a state from which they could not be rescued, even if some individual genes showed signs of reversion. In addition, the expression trend of cellular senescence markers remained relatively unchanged throughout the time course. When we categorized senescence-related genes into inducers and inhibitors, we observed a slight change in the expression of inducers and a gradual decline in inhibitors (Supp. Figure 2A), suggesting that weakened inhibition may ultimately lead to senescence. It is possible that, upon SIRT6 depletion, the cell population harbors a small percentage of senescent cells; however, the system itself is not senescent, as the cells continue to divide throughout the experiment. Overall, the results of this analysis are in correlation with previously published studies and overlap with altered pathways in other aging models and aging tissues^11,20–22,26–28^, thereby strengthening our confidence in this model for aging research.

**Figure 2:**
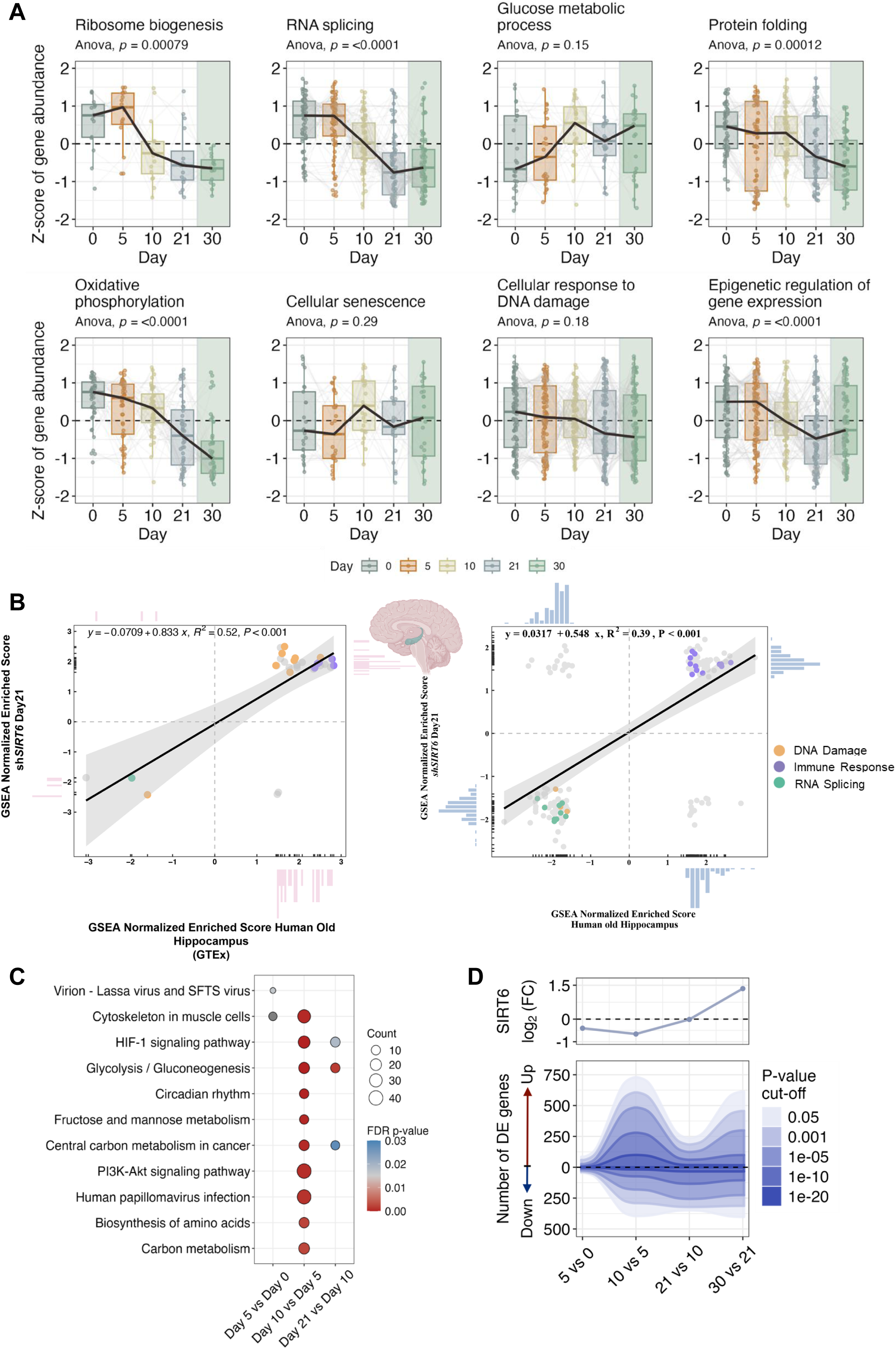
Dynamic Changes in Aging-Related Pathways Under Gradual SIRT6 Decline. **(A)** Plot showing dynamic changes in eight aging-related pathways during SIRT6 silencing (Days 0, 5, 10, and 21) and after recovery (Day 30, green background). **(B)** Correlation plot of gene sets enriched in human aging hippocampus and shSIRT6 Day 21 cells based on GTEx (left) and microarray (right) datasets, with selected pathways highlighted. **(C)** Enrichment analysis of DEGs between consecutive time points. Colors represent FDR-adjusted P values, and circle sizes are proportional to gene counts. **(D)** The number of differentially expressed (DE) genes in consecutive comparisons is shown across multiple P-value cutoffs, with upregulated and downregulated genes quantified separately. The upper panel shows SIRT6 log2 fold change, revealing that the magnitude of transcriptional changes, particularly the number of upregulated genes, correlates with the extent of SIRT6 silencing.

While analyzing known hallmarks of aging provides initial validation for our model, assessing its relevance to tissue-level aging, particularly in the brain, requires direct comparison with transcriptomic data from aged human tissues. To address this, we compared Day 21 transcriptomics with aging-associated gene expression signatures derived from two independent human cohorts: the Genotype-Tissue Expression (GTEx) dataset and a published human brain microarray study^29^. Aging-related DEGs were identified by comparing young (20-40 years old) and old (≥60 years old) individuals in each brain region, followed by Gene Set Enrichment Analysis (GSEA) and correlation analysis with the Day 21 transcriptome (for the microarray-based analysis, a permissive approach using nominal p-values was applied to account for data biases; see Methods). Our results revealed a consistent association between prolonged SIRT6 silencing in the cellular model and human brain aging signatures across all tested brain regions (Suppl. Table 2).

We began by examining multiple cortical regions. Across both human cohorts, cortical regions showed a clear association between aging-related transcriptional programs and the Day 21 state (Suppl. Figure 2B). Next, we examined the hippocampus (HIP), a brain region that exhibits pronounced transcriptional changes during aging and early neurodegeneration and is regulated by SIRT6^18,30^. The hippocampus similarly displayed robust associations, with enrichment of immune-, DNA damage-, and RNA-related pathways, which have previously been linked to SIRT6 regulation^11,14,31–33^. To further examine these aging-associated signatures, we analyzed both immune– and DNA damage–related categories from the GTEx dataset at the individual gene level and found strong correlations with Day 21 (Suppl. Figure 2C), further confirming preservation of aging-associated transcriptional changes in the shSIRT6 model.

Finally, we examined the mediodorsal (MD) thalamus and the hypothalamus, two regions with established functional connectivity. Comparison with Day 21 revealed enrichment of metabolic stress and RNA processing–related pathways in the MD thalamus, whereas the hypothalamus showed association with immune– and extracellular matrix–related pathways (Suppl. Figure 2D, Suppl. Table 2), demonstrating that the shSIRT6 model captures aging-associated alterations in functionally connected brain regions. Overall, comparisons across independent datasets and platforms revealed convergence on shared aging-related transcriptional themes, indicating that Day 21 SIRT6 depletion captures conserved molecular features observed in human brain aging.

Having established that gradual SIRT6 downregulation recapitulates features of human brain aging, we next leveraged this system to investigate how aging unfolds as a gradual process through the lens of SIRT6. We expected to see that SIRT6 levels and their target genes would change gradually and in a structured order. We hypothesize that genes critical for maintaining cell identity and specialized functions, such as neuron-specific transcription factors involved in terminal differentiation, will be deregulated early in the aging process. In contrast, genes essential for cellular maintenance, survival, and metabolism, including pro-apoptotic and housekeeping genes, will be affected later. This hypothesis is supported by studies demonstrating increased variability and deregulation of specialized gene expression during aging, particularly in the brain, which further highlights the structured and hierarchical nature of the aging process^34,35^. To better understand the gradual changes in aging, we conducted an enrichment analysis of DEGs by comparing each pair of consecutive time points (Figure 2C). We found that most changes from Day 10 involved pathways that are not critical for sustaining cell viability, such as circadian rhythms and alterations in metabolic pathways. These processes exhibited fluctuating expression trends, with an initial upregulation by Day 10 followed by downregulation between Day 10 and Day 21. Pathways related to stress responses, such as AMPK signaling, became activated as the process progressed, between Day 21 and Day 30 recovery, suggesting a shift in cellular responses (Suppl. Figure 2E, 2F). In addition, we found that the number of DE genes, particularly the number of upregulated genes, are correlated with the level of SIRT6 silencing (Figure 2D). These results demonstrate how our model can be used to investigate not only the global changes that occur during aging but also their temporal order.

### Revealing Dynamic Transcriptional Changes During SIRT6 Silencing Linked to Aging and Neurodegeneration

To understand aging as a process, the time-dependent changes, and discover novel pathways, we performed a comprehensive analysis of gene clustering and identified 24 distinct gene expression trends during the gradual silencing of SIRT6 (Figure 3A, Suppl. Figure 3A). Each group captures a unique dynamic of gene abundance across the five time points (Day 0, 5, 10, 21, and 30 Recovery), revealing the complexity of transcriptional responses. Interestingly, the expression patterns show diverse behaviors, including oscillating, reversible, and irreversible trends, indicating that these changes are not linear. Instead, they suggest a dynamic process of change and adaptation across several clusters in alignment with recent longitudinal multi-omics analyses across the human lifespan^36^. Further enrichment analysis of these clusters revealed significant overrepresentation of critical biological pathways (Figure 3B). The 15 enriched groups showed pathway enrichment for both expected and novel functions impaired during aging. Clustering by expression dynamics provides higher-resolution separation, allowing us to better distinguish the dynamic and non-linear processes occurring, along with shared pathways, from pathways uniquely enriched in specific clusters. For instance, clusters 2, 12 and 19 share similarities in the central hub of neurodegenerative diseases, but differ by excluding cellular processes like nuclear-cytoplasmic transport and cell cycle. In addition, while all categories found in cluster 12 are also present in cluster 2, cluster 19 introduces a unique focus on reactive oxygen species (FDR p-value = 0.012), demonstrating how expression-based clustering reveals subtle regulatory differences between overlapping pathways. In other groups, additional aging-related pathways are highly represented, including HIF-1 signaling pathway (FDR p-value < 1 × 10⁻⁸) and glycolysis/gluconeogenesis (FDR p-value = 2.1 × 10⁻⁵) in cluster 1. This comprehensive analysis not only delineates the dynamic changes in gene expression associated with SIRT6 silencing, but also demonstrates the capacity of our model to capture the non-linear trajectories observed in aging, as seen in other omics datasets, while highlights the potential to uncover novel insights into the molecular mechanisms of aging and neurodegenerative diseases, the complexity of the dynamic changes, and what changes are more easily reversed.

**Figure 3:**
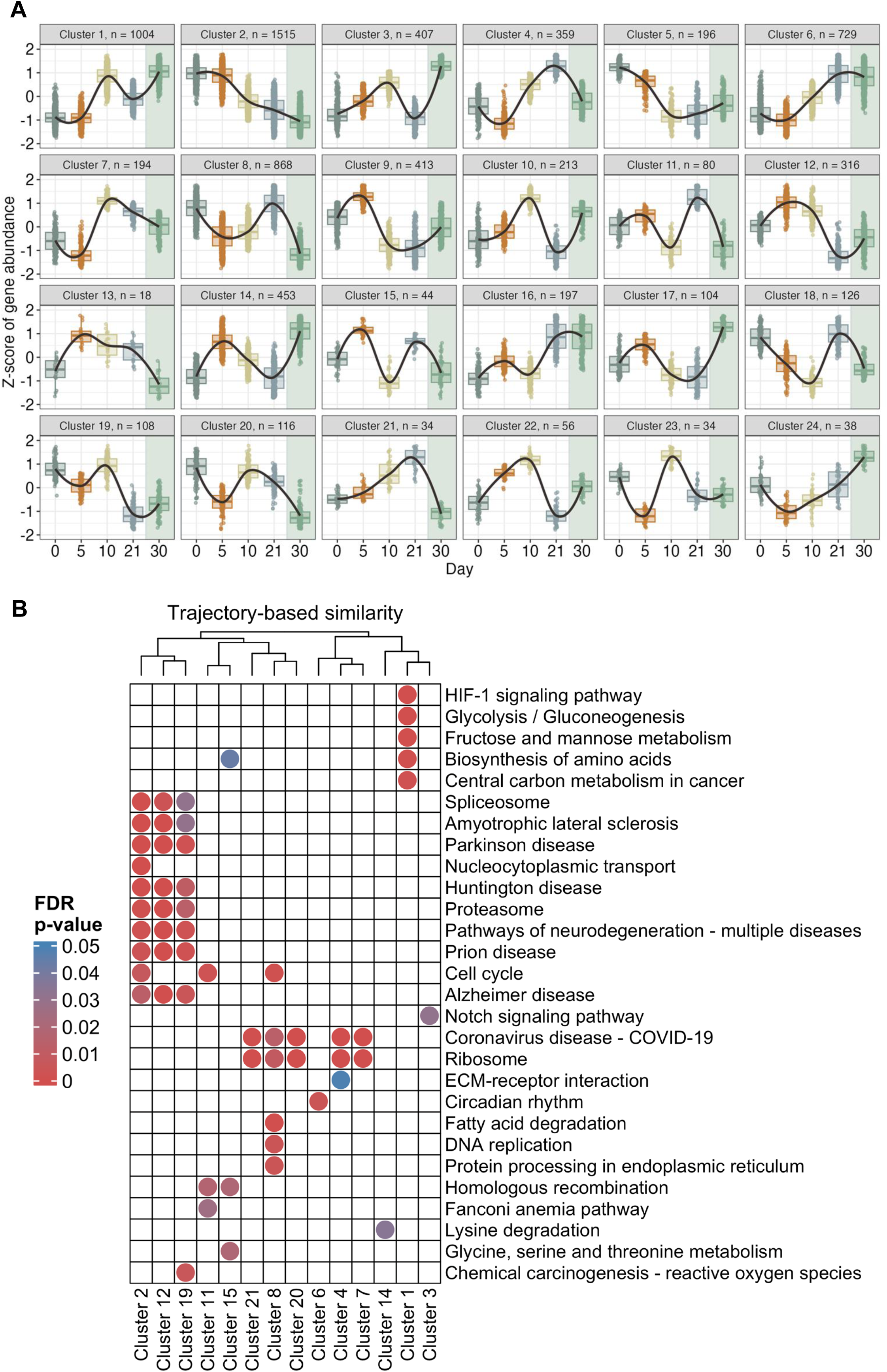
Revealing Dynamic Transcriptional Changes During SIRT6 Silencing Linked to Aging and Neurodegeneration. **(A)** Clustering of 24 gene groups based on expression trends during SIRT6 silencing (Days 0, 5, 10, and 21) and after recovery (Day 30, green background). **(B)** Enrichment analysis of gene clusters, with clusters shown on the x axis and enriched pathways on the y axis. Colors indicate FDR-corrected p values from a hypergeometric test. The column dendrogram was generated by hierarchical clustering based on the genes associated with each cluster shown in the figure.

### Dynamic Gene Clusters Reveal Oscillatory Stress Pathways and Nuclear Decline During SIRT6 Silencing

To better understand some of the clusters, we analyzed enriched terms. In addition, we added a time point with extended SIRT6 silencing for 30 days to more effectively capture the progressive changes, providing a clearer contrast with the recovery condition. We focused on four clusters, which were found to be enriched in key aging indicators, demonstrating trends of consistent downregulation (cluster 2) and oscillatory dynamics (clusters 8, 11 and 15) (Figure 4A). Cluster 2 was enriched for nuclear integrity, while oscillatory clusters were enriched for cellular stress, apoptosis, and genomic instability. Therefore, we measured DNA damage using ɣH2A.X staining, which identified clusters 11 and 15. We observed an increase in ɣH2A.X foci five days post-SIRT6 silencing, followed by a decline at Day 10, and a resurgence from Day 15 to Day 30 (Figure 4B, Suppl. Figure 4A). Notably, the number of cells with more than five foci significantly decreased by Day 30 Recovery, suggesting that SIRT6 reintroduction mitigates DNA damage. We speculated that the decrease at Day 10 may reflect loss of the most damaged cells through cell death, after which the remaining cells begin to accumulate damage again in an oscillatory manner. Therefore, we evaluated cell viability through TUNEL staining (Suppl. Figure 4B). This assessment of apoptotic cell state revealed an increase in apoptotic cells upon SIRT6 silencing, peaking at Day 10 and followed by a sharp decrease by Day 15 (Figure 4C), reflecting the clearance of apoptotic cells from the viable cell population, which increases again when SIRT6 is not re-expressed. These results suggest that while initial apoptosis was cleared, subsequent cell viability was compromised, which may partially explain the observed oscillatory gene clusters. The results show a correlation between DNA damage and apoptotic cell numbers (Supp. Figure 4C). In the early stage of Day 5, DNA damage is driving apoptosis. As apoptotic cells are cleared at Day 10, DNA damage appears low, respectively. However, persistent SIRT6 silencing prevents efficient DNA repair, leading to recurrent DNA damage and increased apoptosis at later time points of Day 15, 21 and 30. This cycle of damage and cell death may explain the oscillatory patterns observed in gene expression clusters. Moreover, SIRT6 reintroduction successfully restores both DNA damage and apoptosis to low levels, highlighting its critical role in maintaining genomic stability and cell survival.

**Figure 4:**
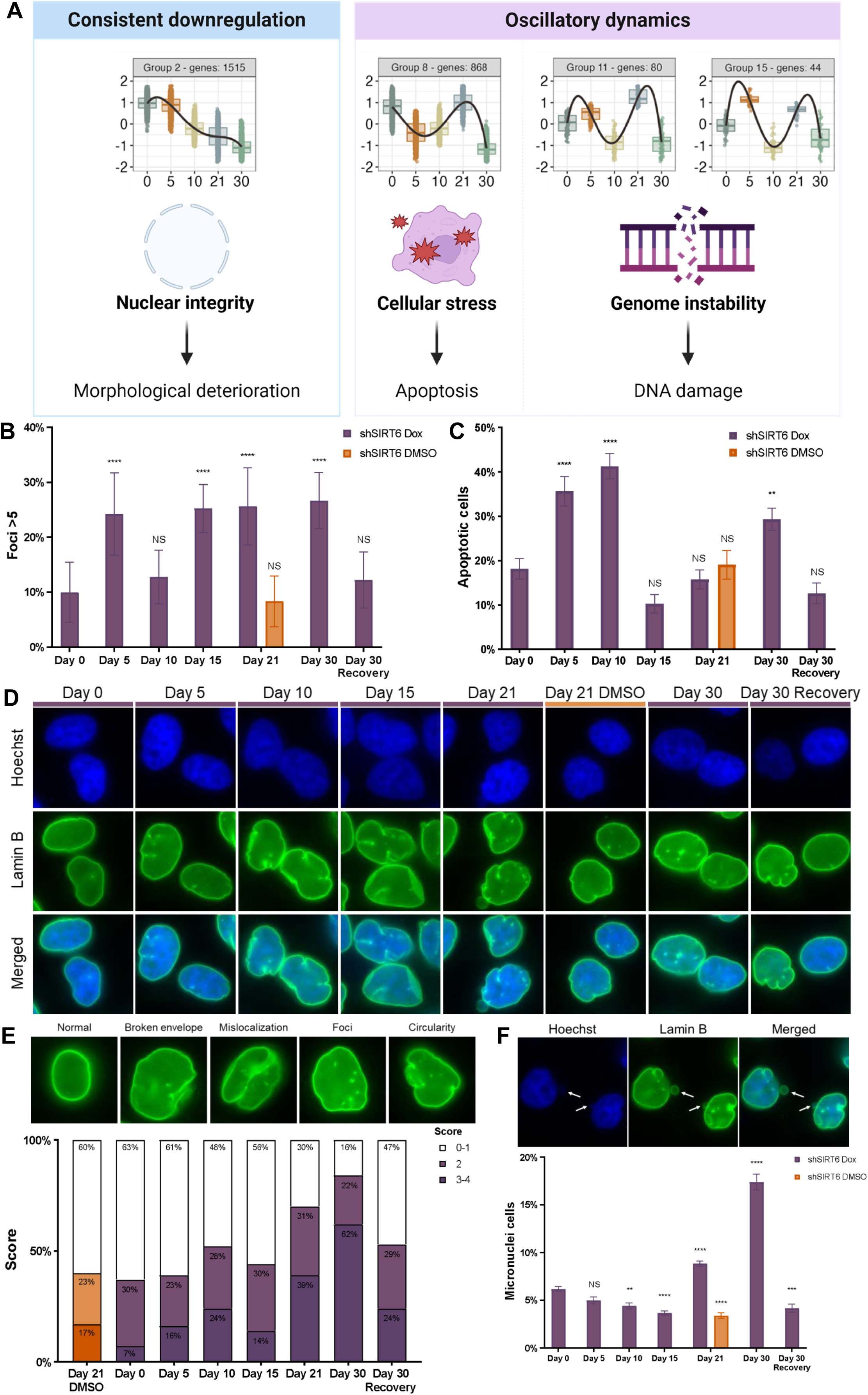
Revealing Dynamic Transcriptional and Cellular Changes During SIRT6 Silencing Linked to Aging and Neurodegeneration. **(A)** Cluster 2 shows consistent downregulation, whereas clusters 8, 11, and 15 display oscillatory expression patterns. These clusters were further analyzed for markers of genomic instability, cellular stress, and nuclear integrity. **(B)** Quantification of the percentage of cells containing five or more γH2A.X foci per nucleus. **(C)** Quantification of apoptotic cells over time. **(D)** Representative immunofluorescence images of Lamin B in cells subjected to SIRT6 silencing. **(E)** Nuclear integrity assay during SIRT6 decline. Upper, images of normal nuclei and four representative defective nuclei illustrating the parameters used to assess nuclear integrity. Lower, summary of four-parameter matrix scoring showing significant differences between observed and expected frequencies determined by a chi-square test (χ² = 129.5, df = 14, P < 0.0001). **(F)** Representative images of cells with micronuclei (upper) and corresponding quantification (lower) during prolonged SIRT6 silencing and recovery. Data are presented as mean ± SEM. (ANOVA; NS, not significant; *P < 0.05, **P < 0.01, ***P < 0.001, ****P < 0.0001).

Given that aging leads to functional decline and morphological deterioration of the nucleus, and that a deformed nucleus contributes to accelerated aging, as seen in laminopathies^37^, we examined nuclear integrity by immunofluorescence of Lamin B (Figure 4D). This protein is crucial for the nuclear lamina and was found in cluster 2, which is enriched for neurodegenerative disease pathways and cell cycle functions. Lamin B was found in cluster 2 alongside other key nuclear integrity genes, including SYNE1, BANF1, and ZMPSTE24, all play crucial roles in maintaining nuclear envelope integrity and proper nuclear architecture, ensuring the structural stability and function of the nucleus. Moreover, we have previously observed a decline of Lamin B and nuclear integrity in cortical neurons from aged mice^38^. We assessed nuclear integrity using four parameters: ‘broken envelope’, ‘mislocalization’, ‘foci’, and ‘circularity’ (Figure 4E, upper panel). A score of zero indicates an intact nucleus, while a score of four reflects maximal damage. As SIRT6 expression decreased, the nuclear structural integrity was compromised. Day 30 Recovery showed moderate improvement in nuclear integrity compared to Day 30 but did not fully return to Day 0 levels (Figure 4E, lower panel). Analysis of each parameter revealed that ‘mislocalization’ and ‘circularity’ were the most affected (Suppl. Figure 4D). While ‘mislocalization’ improved somewhat during recovery, circularity’ and ‘broken envelopes’ did not fully recover, reflecting persistent nuclear deformities that may not be reversible once the envelope is damaged (Suppl. Figure 4E). Moreover, to evaluate genomic instability during prolonged SIRT6 silencing, we quantified the percentage of cells exhibiting micronuclei. From Day 0 to Day 15, 3-6% of the cells displayed micronuclei. By Day 21, the percentage of cells with micronuclei increased to approximately 8%, suggesting an accumulation of genomic damage. The most pronounced rise was observed at Day 30, with 17% of cells exhibiting micronuclei, highlighting a sharp escalation in nuclear abnormalities (Figure 4F). This trend aligns with increasing nuclear structure and integrity scores corresponding to SIRT6 decay^38,39^. Notably, following SIRT6 re-expression, the percentage of cells with micronuclei returned to baseline levels, demonstrating full recovery and underscoring the essential role of SIRT6 in maintaining genomic integrity. The nuclear lamina provides structural support to the nucleus, and its integrity is essential for maintaining proper nuclear dimensions. Therefore, we measured nuclear size in SHSY5Y-SIRT6 wild-type (WT) and knockout (KO) cells. Under regular conditions, we observed no significant differences in nuclear size between WT and KO cells, suggesting a capacity to maintain normal nuclear size even in the absence of SIRT6 (Suppl. Figure 4F, NT), as well as lack of significant senescent population, that tend to have enlarged nuclei However, upon treatment with Leptomycin B (LMB), a nuclear export inhibitor, nuclear size significantly increases in both conditions, with an even more pronounced effect in SIRT6 KO cells (Suppl. Figure 4F, LMB). This finding suggests that although the absence of SIRT6 alone did not significantly affect nuclear size, additional stress led to nuclear enlargement, indicating that SIRT6 plays a protective role under stress conditions in maintaining nuclear structure.

Overall, these results allow us to understand the nature of the oscillation for certain cluster categories, such as DNA repair and apoptosis. Moreover, our system enables us to investigate the reversibility of the phenomena, not only at the transcriptomic level, but also in terms of changes in cellular morphology, apoptosis, nuclear envelope and DNA damage. This comprehensive approach allows us to better identify potential targets for future research, and highlights the advantage of analyzing multiple time points, as oscillatory behaviors may mask significant alterations when looking only at one time point.

### Irreversible Downregulated Genes Reveals Shared Neurodegenerative Pathways and a Distinct Nucleocytoplasmic Transport Pathway

We decided to focus on cluster 2 since it contained several neurodegenerative diseases, denoting changes that are pathological in this cluster. First, we took advantage of our brS6KO mouse model RNA-seq where we have already confirmed a neurodegenerative phenotype with behavioral changes, cell death, and increase in markers for neurodegeneration, such as hyperphosphorylated Tau, and DNA damage^18^. Cluster 2, which included neurodegenerative diseases, was most similar to the top enriched categories in the mouse brains^26^. The overlap confirmed that although we are using a cell line, the changes parallel those observed in the mouse brains (Suppl. Figure 5A).

Next, we focused on all the genes associated with neurodegenerative diseases from cluster 2 to better understand the molecular functions they encompass. Those genes are linked to five major neurodegenerative diseases: Alzheimer’s, Parkinson’s, Huntington’s, ALS, and prion diseases. We then identified the overlapping genes across all five neurodegenerative diseases (NDDs), resulting in a subset of 34 common genes that share a central core with two main molecular functions clusters in oxidative phosphorylation and proteasome (Suppl. Figure 5B), suggesting these are central pathways affected in all neurodegenerative diseases and regulated by SIRT6. To further understand the interconnected pathways, we performed a gene network analysis of the full set of 112 NDDs related genes using KEGG and molecular function annotations. This revealed a central hub NDDs connected to oxidative phosphorylation, as well as to longevity, cellular senescence and apoptosis, suggesting common genes may serve as regulatory nodes that integrate these processes within a broader neurodegenerative landscape (Suppl. Figure 5C). Out of all groups comprising the gene network, nucleocytoplasmic transport (NCT) was especially noticeable since it was separated from all the other categories. Therefore, we decided to focus on this pathway as an independent mechanism that is affected by SIRT6 and common to neurodegenerative diseases. When we examined the genes in cluster 2 associated with NCT, we observed major enrichment in nuclear export, nuclear pore and mRNA transport. This category is particularly interesting since nuclear stability and pore are affected in aging and progeria. In addition, NCT is compromised in certain diseases, such as ALS, all of which exhibit decreased SIRT6 levels, leading us to hypothesize that SIRT6 could be regulating this process.

### SIRT6 Role in Regulating Nucleocytoplasmic Transport During Aging

Disruption in NCT could be caused by several reasons, such as defects in the nuclear pore complex, altered mRNA cargo, or dysregulation of shuttling proteins. While SIRT6 has not previously been associated with NCT, our findings suggest that our model could be used in identifying new, relevant pathways. We therefore sought to investigate the role of SIRT6 in NCT to evaluate our model prediction in impaired processes.

To test whether SIRT6 could regulate NCT, we started by expressing a synthetic NCT system of a dual reporter plasmid^40^ in SIRT6 WT and KO cells. This plasmid consists of a GFP sequence fused to NES sequence (GFP:NES), an internal ribosome entry site (IRES) and RFP sequence fused to an NLS sequence (RFP:NLS) (Figure 5A). To examine disruptions in nuclear export and import, we used selective inhibitors – Leptomycin B (LMB), a specific export inhibitor, and Ivermectin (IVM), a specific import inhibitor (Figure 5B). We tested SIRT6 WT and KO cells under no-treatment conditions and after 24 hours of treatment with LMB and IVM (Figure 5C-D, Suppl. Figure 5D). In WT cells, the response to both inhibitors was as expected, and RFP levels were not significantly different. LMB treatment reduced cytoplasmic GFP levels (Figure 5C, left panel, LMB) and led to a corresponding nuclear accumulation (Figure 5D, left panel, LMB), consistent with its role in blocking nuclear export. IVM treatment, as expected, decreased nuclear GFP levels, reflecting impaired nuclear import. Interestingly, despite being an import inhibitor, IVM also reduced cytoplasmic GFP levels, suggesting a possible indirect effect on nuclear export or translation (Figure 5C-D, left panel, IVM). In contrast, SIRT6 KO cells showed no significant response to either treatment (Figure 5C, right panel), except for an increase in nuclear GFP levels following LMB treatment, indicating export inhibition (Figure 5D, right panel). However, cytoplasmic GFP remained unchanged, unlike in WT cells, where it was reduced. IVM treatment had no effect at all on GFP levels in either the nucleus or the cytoplasm (Figure 5C-D, right panel, IVM). Taken together, these results indicate that SIRT6-KO cells are less responsive to inhibitor treatment, with a more pronounced effect in the cytoplasm. This suggests a basal impairment of the NCT machinery, potentially to the extent that it is already sufficiently compromised to no longer respond to drug treatment.

**Figure 5:**
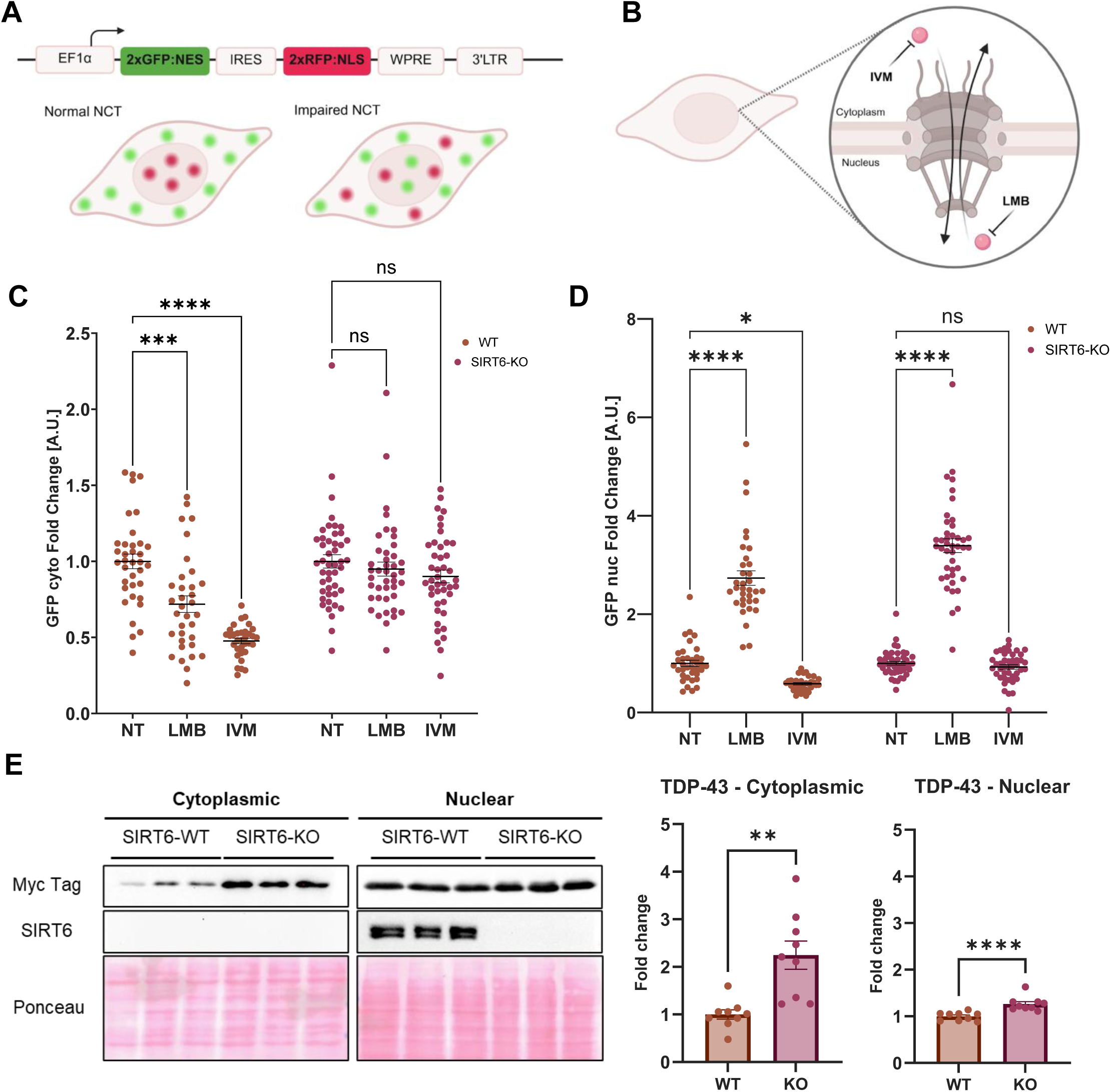
SIRT6 Role in Regulating Nucleocytoplasmic Transport During Aging. **(A)** Diagram of the GFP:NES–i–RFP:NLS lentiviral reporter used for expression in SIRT6 wild-type (WT) and knockout (KO) cells. **(B)** Schematic illustration of ivermectin (IVM), a specific nuclear import inhibitor, and leptomycin B (LMB), a specific nuclear export inhibitor. **(C,D)** Quantification of GFP signal in the cytoplasm (C) and nucleus (D) of SIRT6 WT and KO cells under basal conditions and after 24 h treatment with LMB or IVM (ANOVA). **(E)** Representative western blot analysis of cytoplasmic and nuclear fractions from SIRT6 WT and KO SH-SY5Y cells. Left, representative blots; right, quantification of Myc-tagged TDP-43 levels normalized to Ponceau total protein staining (Welch’s t-test for cytoplasmic TDP-43; Mann–Whitney test for nuclear TDP-43). NS, not significant; *P < 0.05, **P < 0.01, ***P < 0.001, ****P < 0.0001.

To assess our results in a more physiologically relevant protein, we next examined TDP-43, an RNA-binding protein with a crucial role in NCT, which is disrupted in neurodegenerative diseases, including ALS, where NCT dysfunction has been identified as a central disease mechanism^41^. We overexpressed TDP-43 in SIRT6 WT and KO cells and found that their levels increased in the cytoplasm and nucleus of KO cells, but to different extents (Figure 5E). While the nuclear fraction showed a modest ∼1.2-fold increase compared to WT, the cytoplasmic levels of TDP-43 were elevated by more than twofold; this cytoplasmic TDP-43 has been associated with ALS. Moreover, we observed greater variability in the cytoplasmic fraction across replicates, possibly indicating a more dynamic, context-dependent regulation in the cytoplasm rather than a more robust nuclear phenotype. These findings suggest that SIRT6 plays a role in regulating general protein transport machinery (as shown by the GFP:NES-RFP:NLS system), but also the transport of important specific proteins in neurodegeneration, and illustrate how our model can uncover critical aging-related pathways regulated by SIRT6. Further investigation into the interplay between SIRT6 and nucleocytoplasmic transport machinery may provide mechanistic insights into how SIRT6 depletion disrupts cellular compartmentalization during aging and disease.

### Reversible and Irreversible Changes in Gene Expression Following SIRT6 Silencing and Recovery

One of the main advantages of working with cell lines, compared to animal models or tissues, is the ease with which they can be manipulated. In our cellular system, SIRT6 can be readily re-expressed by simply removing Dox from the cell medium, allowing us to distinguish between reversible and irreversible changes in gene expression and phenotypes associated with SIRT6 silencing. To better understand these changes, we conducted additional independent RNA-seq analyses of two experimental conditions: continuous 30-day Dox treatment without recovery (Day 30) and a 21-day Dox treatment followed by a 9-day recovery period (Day 30 Recovery) (Suppl. Figure 6A). Comparing these two time points with Day 0 helps distinguish processes that can be rescued by SIRT6 reintroduction from those that remain irreversibly altered. As expected, differential expression analysis revealed a greater number of DEGs after 30 days of continuous SIRT6 silencing than after the 9-day recovery period (Figure 6A, Suppl. Figure 6B). This reduction in DE genes upon recovery suggests that reintroducing SIRT6 mitigates some of the gene expression changes induced by prolonged silencing.

**Figure 6:**
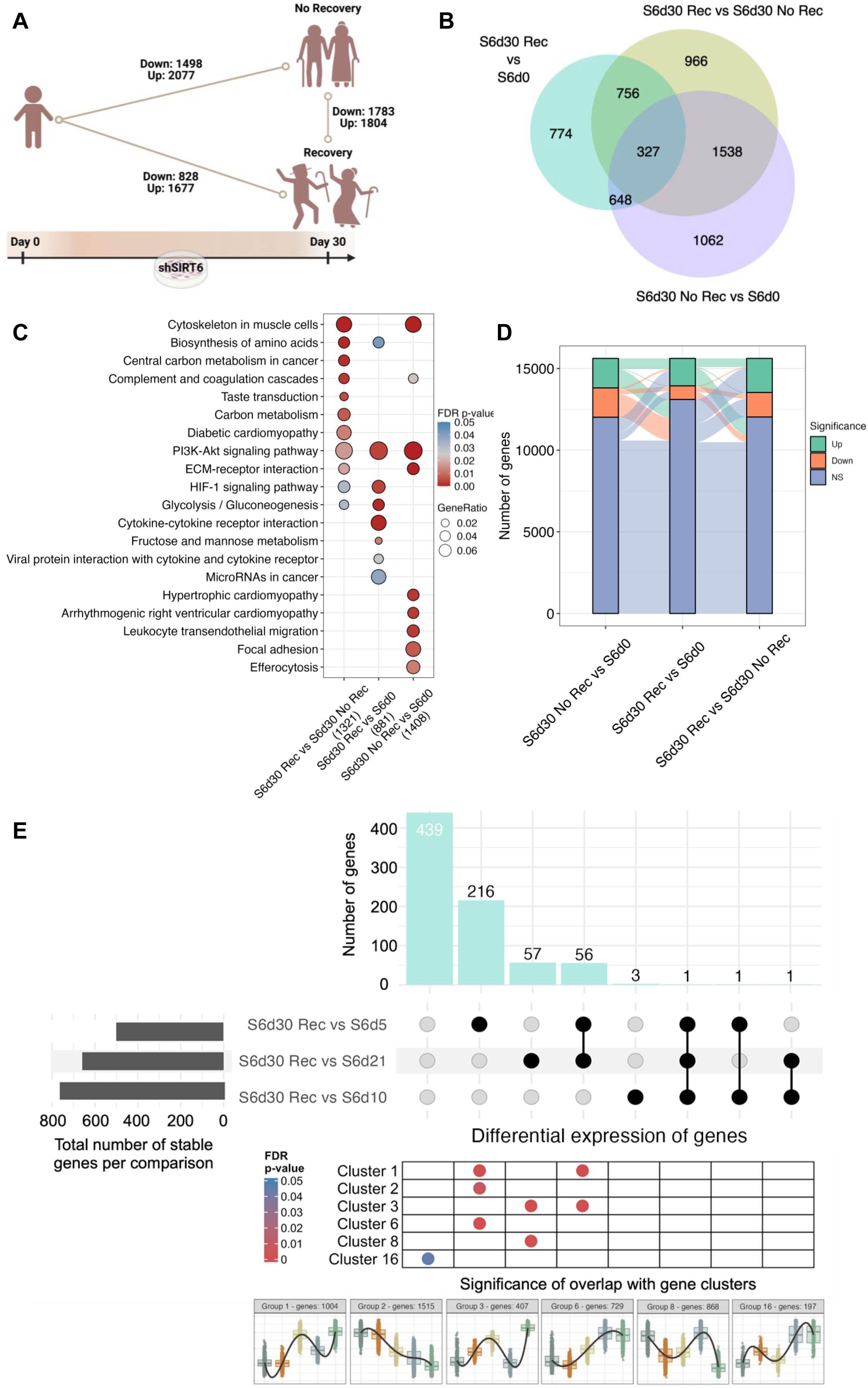
Reversible and Irreversible Changes in Gene Expression Following SIRT6 Silencing and Recovery. **(A)** Schematic illustration of the three experimental conditions (Day 0, Day 30 without recovery, and Day 30 recovery) and the number of differentially expressed genes (DEGs) identified between them. **(B)** Venn diagram showing shared and condition-specific DEGs across the three experimental conditions. **(C)** KEGG pathway enrichment analysis of the union of DEGs within each comparison group (Day 0 vs. Day 30, Day 0 vs. Day 30 recovery, and Day 30 vs. Day 30 recovery), as defined by the full circles in the Venn diagram. **(D)** Alluvial plot illustrating transitions of upregulated, downregulated, and non-significant (NS) genes across the three comparisons. **(E)** UpSet plot showing pairwise comparisons between Day 30 recovery and Day 5, Day 10, and Day 21. Horizontal gray bars indicate the total number of stable (non-DE) genes in each comparison. Vertical light turquoise bars indicate the number of overlapping genes between the 774 unique Day 30 recovery DEGs and each earlier time point. Black dots indicate inclusion in a given comparison; gray dots indicate exclusion. The heatmap at the bottom shows the significance of overlap (hypergeometric test; only overlaps with FDR-adjusted p values < 0.05 are shown) between these genes and the gene clusters defined in Fig. 3A.

To further elucidate the impact of SIRT6 silencing and recovery on gene expression, we examined three key comparisons: Day 0 vs. Day 30, Day 0 vs. Day 30 Recovery, and Day 30 vs. Day 30 Recovery (Figure 6B). These comparisons respectively capture the cumulative transcriptomic changes following 30 days of SIRT6 depletion, the changes present after recovery, and the specific transcriptional response to SIRT6 re-expression. To functionally interpret these gene expression changes, we performed KEGG enrichment analysis of the union of DEGs within each comparison group (Day 0 vs. Day 30, Day 0 vs. Day 30 Recovery, and Day 30 vs. Day 30 Recovery) (Figure 6C). In the Day 0 vs. Day 30 group, enriched pathways included immune-related processes reflecting stress-associated responses commonly observed during aging. Unique genes in this group were enriched in key aging-related pathways, including ribosome biogenesis, associated with proteostasis, and base excision repair, a central pathway in DNA damage repair (Suppl. Figure 6C). These findings further support the role of this group in capturing transcriptomic signatures characteristic of aging. In the Day 0 vs. Day 30 Recovery comparison, enriched pathways included glycolysis/gluconeogenesis and amino acid biosynthesis, among others, reflecting adaptive transcriptional programs specifically activated by SIRT6 re-expression and may support metabolic recovery and stress adaptation. For the Day 30 vs. Day 30 Recovery comparison, the changes depended on active SIRT6 function, since they were not differentially expressed when compared to Day 0. We observed significant enrichment for metabolic pathways, including amino acid biosynthesis and glycolysis/gluconeogenesis, consistent with SIRT6’s known role in regulating these processes ^9^ (Figure 6C). Notably, KEGG analysis of unique genes in this comparison revealed enrichment of neurodegeneration-related pathways, including Huntington’s disease, prion disease, and Parkinson’s disease (Suppl. Figure 6C). These findings suggest that SIRT6 re-expression reactivates genes involved in neuronal maintenance and may identify disease-relevant reversible targets. To further dissect the dynamics of gene expression reversibility, we track the transitions of genes categorized as upregulated, downregulated, or not significant (NS) across the three comparisons (Figure 6D). This analysis revealed only a small fraction of genes that switched directly between up– and downregulation, whereas a much larger proportion shifted from being up– or downregulated to NS after SIRT6 re-expression. Notably, many genes that were differentially expressed at Day 0 vs. Day 30 became NS in the recovery condition, underscoring the rescue potential of restored SIRT6 activity. Moreover, many of these same genes reappeared as upregulated or downregulated in the Day 30 vs. Day 30 Recovery comparison, while others transitioned from being differentially expressed at Day 0 vs. Day 30 Recovery to NS at Day 30 vs. Day 30 Recovery. Together, these patterns suggest that the expression of these genes is directly modulated by active SIRT6 function, further supporting the model’s capacity to capture both reversible and persistent aspects of aging-associated transcriptomic changes.

Based on our transcriptomic analysis comparing the prolonged 30-day SIRT6 silencing condition with the recovery phase, we thought that SIRT6 re-expression might only partially reverse the effects, not fully restore the Day 0 state. To explore the potential stage at which recovery-related pathways become most prominent, we compared the Day 30 Recovery profile with each of Day 5, Day 10, and Day 21 time points. This comparison allowed us to determine which time point’s gene expression pattern most closely resembles the recovery state. The comparison revealed that the highest number of stable genes – defined as genes not significantly differentially expressed – was observed between Day 10 and Day 30 Recovery, indicating that Day 10 is transcriptionally most similar to the recovery state compared to Days 5 and 21 (Figure 6E, gray bars). To examine this further, we focused on the 774 unique DEGs identified in the comparison between Day 0 and Day 30 Recovery (Figure 6B). We found substantially greater overlap with Days 5 and 21, whereas only minimal overlap was observed with Day 10 (only three overlapping genes). (Figure 6E, turquoise bars). This finding reinforces the notion that Day 10 is the least transcriptionally divergent from the Day 30 Recovery state. Our results suggests that while some aspects of the phenotype can be reversed, the system does not fully return to a “young” state but rather stabilizes at an intermediate, more “middle-aged” transcriptional profile, further strengthening the hypothesis that Day 10 represents a critical point when considering recovery-related pathways. Notably, we found that 439 out of the 774 DEGs were unique to the Day 0 vs. Day 30 Recovery comparison, suggesting that these genes may represent transcriptional changes that emerge only after prolonged SIRT6 silencing and are modulated following its re-expression. In addition, we compared the same set of 774 genes with the gene clusters defined in Figure 3A and found that different groups of comparisons showed significant overlaps with distinct clusters – including Clusters 1, 2, 3, 6, 8, and 16 – each of which represents a unique expression pattern and trajectory (Figure 6E, heatmap). These findings further underscore the complexity of modeling aging, as it does not follow a single linear path but instead reflects multiple dynamic trajectories that must be considered when interpreting the potential for recovery. Importantly the recover pathways at the two end-points include mainly metabolic pathways, suggesting recovered cells returned to their normal metabolic use. Among the categories that continue to be enriched after recovery include many pathways involved in stress sensing, integrin mediated signaling and cytoskeleton rearrangement, suggesting extracellular signaling adaptation was not reversed, or slower in its adaptation.

Finally, we found 327 DEGs overlapping across the three experimental conditions (e.g. Day 0, Day 30, and Day 30 Recovery) (Figure 6B). Given the differences in the time points and conditions, we speculated that these common genes exhibited directional differentiation in expression. As we expected, heatmap analysis of these shared genes revealed four distinct subclusters with directionally different expressions (Suppl. Figure 6D), suggesting that although some DEGs are shared across conditions, their behavior, timing, and functional consequences differ, ultimately leading to differences in pathway enrichments and significance levels.

Overall, by taking advantage of our cellular model, we were able to perform precise comparisons between distinct time points and conditions of SIRT6 expression. These results provide insights into the direct effects of SIRT6 and those initiated by its influence on other proteins, distinguishing between transcriptomic changes that are reversible upon SIRT6 reintroduction and those that remain persistently altered. Moreover, they reveal how restoring SIRT6 expression can modulate ongoing changes within affected pathways, highlighting potential rescue interventions. Together, these results deepen our understanding of SIRT6’s role in regulating cellular aging and potential reversibility of age-related changes.

### Gradual SIRT6 Decline Drives Transcriptomic Shifts Toward Pathological Aging and AD

Given the important role of SIRT6 in aging and neurodegeneration, we examined whether the transcriptomic changes in our model reflect those observed in pathological brain aging, with a particular focus on AD. To address this, we compared the RNA-seq data from the SIRT6 knockdown model (Day 10, 21, 30, and 30 Recovery) with published transcriptomes of middle-aged and AD temporal lobe samples^42^. First, by comparing the expression profiles of middle-aged and AD brain samples, we identified DEGs that may be associated with the transition from normal aging to AD (Suppl. Figure 7A). We then performed GSEA to assess the correlation between these AD-related DEGs and the expression profiles at each time point in our model (Figure 7A-D). Finally, to better understand the biological processes underlying these correlations and the shift toward pathological aging mediated by SIRT6, we examined the GO and KEGG enrichment of upregulated and downregulated genes at each time point, focusing on both unique and overlapping pathways (Figure 7E, the full list is provided in Suppl. Table 7)

**Figure 7.**
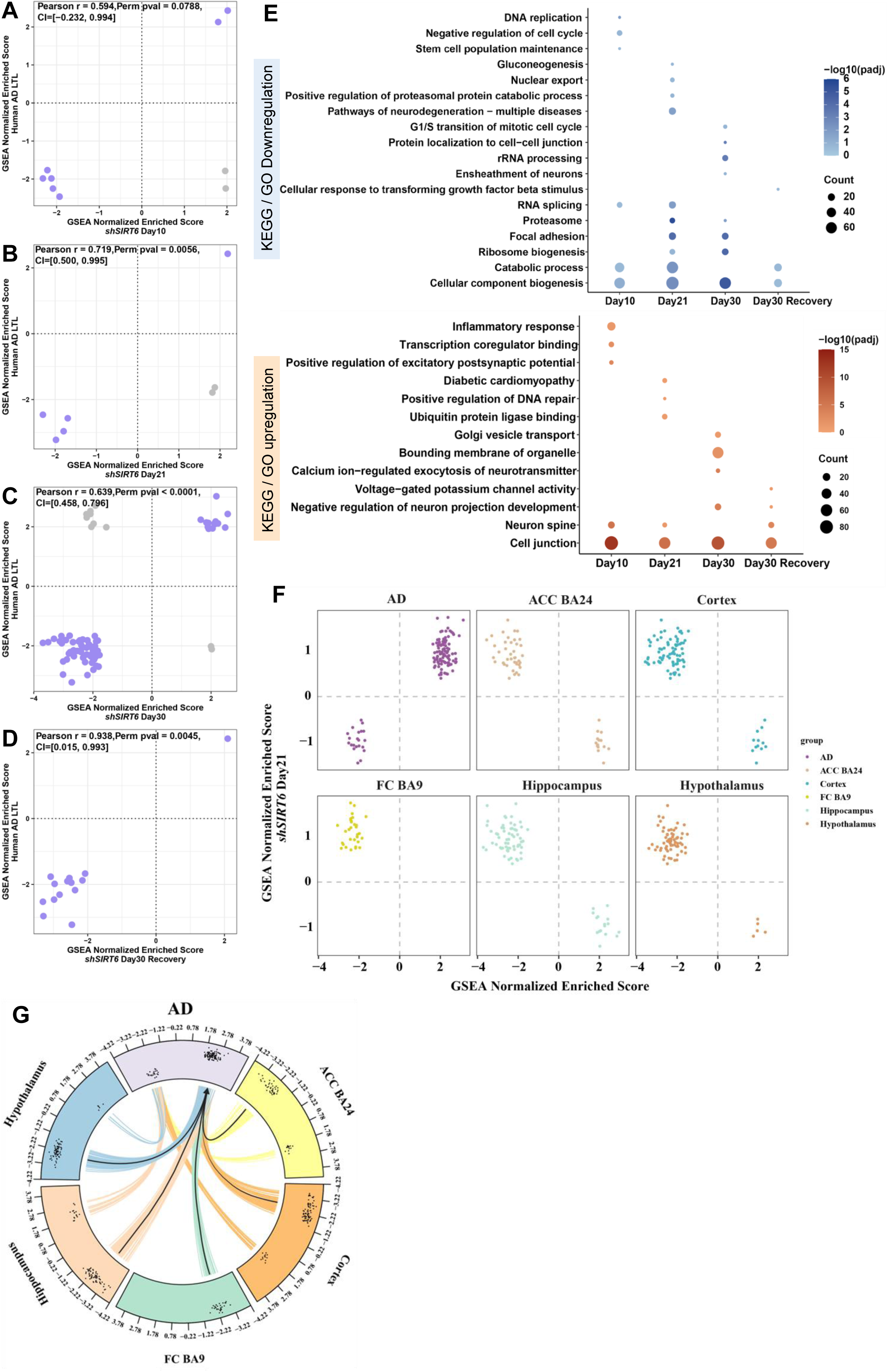
– Gradual SIRT6 Decline Drives Transcriptomic Shifts Toward Pathological Aging and AD. **(A–D)** Correlation plots comparing gene sets enriched in human Alzheimer’s disease (AD) samples with those enriched in shSIRT6 cells at Day 10 (A), Day 21 (B), Day 30 (C), and Day 30 recovery (D). **(E)** Selected enriched Gene Ontology (GO) and KEGG terms shared between human AD samples and shSIRT6 time points. **(F)** Scatter plots showing the distribution of enriched pathways across quadrants based on GSEA normalized enrichment scores in shSIRT6 Day 21 cells (y axis) versus human AD or aging brain regions (x axis). Pathways in AD are concentrated in the first and third quadrants, whereas pathways in aging brain regions are concentrated in the second and fourth quadrants, reflecting opposite enrichment directions. **(G)** Circos plot illustrating pathways that are anti-correlated with aging across multiple brain regions and become positively correlated in AD brains; black arrows indicate pathways showing this transition. Pathways were selected based on Day 30 (padj < 1), AD (padj < 0.05), and aging (padj < 0.05) criteria.

To do so, we implemented a combination of Pearson correlation, permutation testing, and bootstrapping. This approach allowed us to measure the linear relationship between SIRT6 silencing at different time points and the transcriptomic changes observed between adults and AD patients, while also providing a measure of statistical confidence. Our analysis shows that on Day 10, there is no significant correlation between AD and our model, and very few pathways are shared. As the duration of silencing increases, the correlation at Day 21 rises significantly, but with a wide confidence interval (CI) and a small number of correlated pathways. It is only at Day 30 that the correlation, although slightly smaller than on Day 21, exhibits a narrower CI and a very low p-value, along with a significant increase in the number of correlated pathways. These findings suggest a more robust and stable correlation at this time point. Finally, at Day 30 recovery, the number of pathways and confidence decreased significantly, indicating a partial recovery from the SIRT6 knockdown.

#### Day 10 represents early pre-pathological state

At Day 10, despite the highest DEG overlap with AD, the association with AD transcriptome was minimal (Zr=1.67, combining the Pearson’s r and its precision), and there was no significant perm p-value, and very few correlated pathways were present (Figure 7A), suggesting that the overall transcriptional program is still distinct from that of the diseased brain. Functionally, the correlation between Day 10 and AD points to early signs of instability in key cellular maintenance mechanisms that are tightly regulated by SIRT6, including DNA replication and stem cell population maintenance, alongside the upregulation of inflammatory processes (Figure 7E, Day 10).

#### Day 21 represents a transition toward pathological aging

At Day 21, although the association (Zr=1.81, combining Pearson’s r and its precision) with AD transcriptome remains low (Figure 7B), the enriched pathways begin to reflect more pronounced hallmarks of aging controlled by SIRT6. Notably, we observe downregulation of nuclear export, RNA splicing, and synapse organization – indicating emerging metabolic and neuronal dysfunction. In parallel, genes involved in DNA repair and cellular homeostasis – two fundamental processes tightly regulated by SIRT6 – show increased expression (Figure 7E, Day 21). However, it seems to correlate more with aging than with AD, as very few correlated pathways are present (Figure 7B).

#### Day 30 captures core transcriptomic changes of AD

At Day 30, the association with AD transcriptome was the strongest (Zr=6.67, combining the Pearson’s r and its precision) (Figure 7C) and it exhibits the greatest number of overlapping pathways with AD and the strongest overall significance and number of correlated pathways, suggesting at this time point there is a transition from normal aging to pathological aging, and the system becomes more similar to AD (Figure7C). Functionally, Day 30 appears to better capture the transcriptional shifts characteristic of AD (Figure 7E, Day 30). This shift is reflected in the enrichment of several key pathways involved in the immune system, consistent with the neuroinflammatory component of AD, which may reflect activation of glial cells, known to be upregulated in the disease^43^. In parallel, since cell cycle capacity is lost in glial cells and post-mitotic neurons, upregulation of the G1/S transition may indicate inappropriate cell cycle re-entry, as seen in AD where neurons replicate DNA and enter G2 phase^44^. The involvement of glial cells in the disease is also reflected in the enrichment of neuronal ensheathment, including myelination and axon–glial interactions, suggesting altered structural support and communication within the nervous system.

Finally, the downregulation of rRNA processing fits with the broader pattern observed in our model and aging brain datasets, in which RNA metabolism becomes progressively compromised (Figure 7E, Day 30). Together, Day 30 transcriptomic changes align with the AD signature, reinforcing the role of SIRT6 loss in driving pathological aging.

#### SIRT6 re-expression partially reverses AD transcriptomic signatures

Following re-expression of SIRT6 at Day 30 Recovery, the association with AD (R = 5.70, combining Pearson’s r and its precision) dropped to a much lower significance, and the number of correlated pathways indicated a partial reversal of the transcriptome (Figure 7D). Downregulation of the transforming growth factor beta (TGF-β) pathway suggests that the chronic neuroinflammatory response is at least partially mediated by SIRT6, and that its re-expression leads to partial suppression of this pathway. As for neuronal function, the upregulation of voltage-gated potassium channel activity suggests a shift toward rebalancing neuronal excitability and synaptic signaling (Figure 7E, Day 30 Rec). These results once again demonstrate that some transcriptional and functional alterations induced by SIRT6 loss are reversible, reinforcing SIRT6’s therapeutic potential in preventing or delaying pathological aging, and that our model can capture the aging process and the shift toward pathological aging.

Several pathways were found to be enriched across multiple time points, suggesting that they represent core, dynamically regulated processes along the trajectory from early SIRT6 depletion toward pathological aging and, in some cases, into recovery. Changes in their statistical significance over time reflect evolving pathway activity, pointing to both progressive deterioration and potential compensatory responses. Among the downregulated pathways, RNA splicing was enriched at both Day 10 and Day 21, with increased significance at the later time point, consistent with its progressive deterioration observed in our model and its known impairment in AD. Proteasome-related pathways were enriched at Day 21 and Day 30, with stronger enrichment at Day 30, whereas ribosome biogenesis, also present at both time points, showed reduced significance at Day 30. Cellular component biogenesis was consistently downregulated across all time points, with enrichment becoming more significant from Day 10 to Day 30 and decreasing upon recovery. These coordinated changes in RNA processing, protein degradation, and biosynthetic capacity are regulated by SIRT6 and are disrupted in AD, supporting their relevance in the transition from early SIRT6 depletion toward a more pathological aging-like state.

Among the upregulated pathways, negative regulation of neuron projection development was shared between Day 30 and Day 30 Recovery, with stronger enrichment at the Recovery stage. Neuron spine-related pathways were enriched on Days 10, 21, and 30 of Recovery, with the most significant enrichment observed on Day 21. Cell junction pathways were consistently enriched across all time points, with fluctuating significance. These changes reflect dynamic alterations in neuronal structure and connectivity, features commonly affected in neurodegeneration, and further support the progression toward a pathological aging-like state in response to prolonged SIRT6 loss.

Together, these analyses reveal a progressive transcriptomic shift toward an AD-like state during SIRT6 depletion, with partial restoration following its re-expression.

### Opposing Pathway Dynamics in Brain Aging and Neurodegeneration

Having demonstrated that SIRT6 depletion recapitulates core molecular hallmarks of aging and shows a strong association with transcriptomes from healthy aged human brains (Figure 2), we further showed that our model aligns with Alzheimer’s disease (AD) signatures at later time points (Figure 7A-E). However, healthy brain aging and neurodegeneration represent two distinct molecular trajectories. As shown in Fig. 2, comparison with human brain datasets indicated that the model’s Day 21 state closely resembles an aging-associated transcriptional profile. We therefore examined these aging-enriched pathways in the context of AD and observed an anti-correlated distribution (Figure 7F), with pathways clustering in opposing quadrants between aging and AD.

This shift was observed across all tested brain regions, with the majority of transitions involving pathways that were anti-correlated in the Old condition but became correlated in the AD condition (Figure 7G). To conduct this analysis, we focused on identifying pathways undergoing this transition. We used a loose threshold of padj Day30 < 1 to include all pathways from our model, while applying a strict threshold of padj < 0.05 to the Old and AD conditions. This filtering strategy allowed us to specifically identify a high-confidence set of pathways that exhibit a robust and statistically significant change in correlation direction, moving from anti-correlated to correlated (or vice-versa), thereby highlighting the key drivers of this transition. These findings suggest the presence of defined molecular signatures that mark the transition from physiological to pathological aging. Further investigation of these pathways may help identify potential biomarkers that distinguish between normal aging and AD, as well as pathways whose early dysregulation could be reversible. The ability to detect such changes underscores the value of our model in identifying biomarkers and targets for interventions to prevent the shift toward neurodegeneration.

Overall, our results position SIRT6 as a central regulator of brain aging, with its gradual depletion mimicking transcriptomic changes observed across both normal aging and AD. A key advantage of our cellular system lies in its reversibility – SIRT6 can be re-expressed in a controlled manner, enabling the identification of pathways that are modifiable even at late stages. This tractable model offers a valuable platform for aging research, bridging the gap between mechanistic cellular studies and more complex animal or human tissue models. It enables hypothesis generation, early pathway screening, and investigation of temporal dynamics, making it a powerful tool for studying aging and its progression toward neurodegenerative disease.

## Discussion

Current cellular models of aging predominantly rely on senescence^44–46^, a terminal state that has failed to capture aging as a process, pointing to the need for systems that reflect its progressive and dynamic nature, particularly in complex tissues like the brain. SIRT6 has emerged as a central regulator of aging, with its loss driving age-related phenotypes in multiple organisms^12,15,18,23^. Our inducible system offers a distinct advantage by enabling gradual SIRT6 silencing, mimicking aspects of natural aging without inducing senescence. This model provides a powerful tool to dissect early, reversible, and progressive aging-related changes.

The strong transcriptional similarity between our SIRT6 silencing model and aged human brain tissue across independent transcriptomic platforms reinforces the central role of SIRT6 in aging and suggests that its gradual decline recapitulates key molecular hallmarks of brain aging, which are strongly implicated in neurodegeneration. The observed correlations extend beyond global transcriptomic similarity and are reflected at the pathway level, underscoring the biological relevance of the model. The pathways associated with SIRT6 depletion are enriched for processes that are both brain-specific and tightly linked to aging. Notably, many of these pathways are known to be regulated by SIRT6, including DNA damage repair, RNA metabolism and immune system response. SIRT6 is known to suppress inflammation by negatively regulating NF-κB signaling, a major driver of age-related neuroinflammation^47^, and its loss has been associated with elevated expression of pro-inflammatory cytokines, potentially explaining the sustained immune activation observed in our system. The consistent upregulation of immune-related pathways observed under SIRT6 depletion suggests the induction of a chronic inflammatory state that may contribute to neuronal dysfunction and synaptic decline, both characteristic features of cognitive aging and AD^48^. In parallel, the dysregulation of RNA processing and splicing observed in our model may contribute to broader disruptions in neuronal function, including impaired synaptic signaling and cytoskeletal dynamics. The use of a gradual, inducible SIRT6 depletion model, rather than complete loss of function, enhances physiological relevance by more closely reflecting the age-associated decline in SIRT6 activity. In this context, SIRT6 emerges as a central regulator at the intersection of genome maintenance, immune regulation, and brain aging, linking cellular aging mechanisms to region-specific brain vulnerability. The model’s ability to capture conserved SIRT6-regulated aging pathways supports its robustness and relevance as a system for studying aging-associated molecular changes.

To further understand how aging progresses in the absence of SIRT6, we examined the temporal order of transcriptomic changes in our model. We found that early alterations are not essential for cell viability and regulatory processes, such as circadian rhythm and metabolism, while core stress-response pathways like AMPK signaling become activated at later stages. These findings suggest that aging proceeds in a structured manner, not through a specific pathway, but rather prioritizes essential pathways and keeps them regulated, while “neglecting” specialized and less-essential functions, with SIRT6 playing a key role in maintaining the balance between specialized functions and stress adaptation.

A major strength of our model lies in its ability to capture the dynamic and non-linear nature of aging-related gene expression changes. By analyzing multiple time points during the gradual decline of SIRT6, we took advantage of this temporal resolution to uncover a wide spectrum of expression trends, including oscillatory, reversible, and irreversible behaviors that offer a deeper understanding of aging as an ongoing and structured process. The temporal alignment between DNA damage accumulation and apoptosis suggests a cycle in which damaged cells undergo programmed cell death, temporarily clearing the population before damage re-accumulates under persistent SIRT6 silencing. This pattern may explain the oscillatory expression dynamics observed in some gene clusters, with oscillations in other processes suggesting cycles of adaptation. Moreover, our model successfully captures early molecular features of aging and neurodegeneration, such as compromised nuclear integrity^49,50^ demonstrated by alterations in Lamin B.

Compromised nuclear architecture may also contribute to impaired NCT, a critical process mediated by nuclear pore complexes (NPCs) and that tightly depends on their proper function^51,52^. The dysfunction of both NPCs and NCT has been increasingly recognized as a hallmark of aging and a contributing factor in the development of neurodegenerative diseases, including AD and ALS^53,54^. Our model enabled us to identify a previously unrecognized role for SIRT6 in regulating NCT, as evidenced by altered behavior of the physiologically relevant TDP-43 protein in SIRT6-KO cells. Cytoplasmic accumulation of TDP-43 is a hallmark of ALS^55^, one of the multiple neurodegenerative diseases enriched in cluster 2, along with the NCT pathway. TDP-43 is also linked to tauopathy models, where it tends to aggregate in the cytoplasm of neurons^56^. Moreover, Tau, a key protein in several neurodegenerative disorders, is accumulating in the nucleus in the absence of SIRT6, leading to a toxic state for the cell^19^. TDP-43 pathology is defined by a combination of its cytoplasmic aggregation and nuclear clearance, and it is a hallmark observed in 30%-60% of AD cases^57–59^. Moreover, an acetylation switch at a specific residue in TDP-43 RNA-binding domain has been shown to control its function and aggregation^60^. Whether SIRT6 directly deacetylates TDP-43 and whether NCT impairment drives its cytoplasmic accumulation or excessive nuclear clearance remain open questions for further investigation. However, these findings highlight the potential role of SIRT6 in regulating NCT, possibly by influencing NPC function. Importantly, NCT pathway represents only one example of a broader set of aging-associated programs uncovered by our model. The additional pathways identified in our analyses highlight the expansive discovery potential of the model and open multiple directions for future mechanistic investigation into molecular aging.

While SIRT6 re-expression partially restored nuclear structure and reduced DNA damage and apoptosis, certain features, such as nuclear circularity, remained impaired, suggesting that some nuclear defects may be irreversible once structural damage occurs. These findings emphasize the value of the Day 30 Recovery time point, which provides a unique opportunity to distinguish reversible from irreversible aging phenotypes. This stage offers critical insight into processes that may be rescued by restoring SIRT6 function and highlights candidate pathways for therapeutic intervention aimed at halting or reversing cellular decline before it becomes pathological.

Our rescue analysis highlights the strength of using a controlled cellular model to study the dynamics of aging-associated gene expression. Our transcriptomic analysis, incorporating a continuous 30-day SIRT6 silencing condition, allowed us to differentiate between pathways that are persistently altered and those that are potentially reversible. The identification of both reversible and persistent transcriptomic changes provides insight into the temporal regulation of aging pathways and suggests that some aging-like alterations may be mitigated if SIRT6 function is restored early enough. The conditional enrichment analysis suggests that some processes are directly dependent on SIRT6 expression and may be responsive to its restoration. Moreover, our findings position Day 10 as a potential “tipping point” in the trajectory of SIRT6 depletion. At this stage, the cells are already substantially affected by SIRT6 loss, yet they do not fully commit to a fixed aging-associated expression pattern. Instead, they appear to retain some plasticity, with gene expression profiles that partially align with pathways later reactivated during recovery. The ability to reverse gene expression, cellular function, and phenotypes associated with critical cellular aging pathways emphasizes the therapeutic potential of targeting SIRT6-regulated mechanisms to restore cellular homeostasis and delay or prevent pathological aging trajectories.

We further extended our transcriptomic analysis by comparing it with AD brain samples, which highlights a progressive alignment between the SIRT6 depletion model and pathological aging. Although early time points exhibited substantial overlap in DEGs, the expression patterns remained distinct from those of AD, suggesting an early pre-pathological state. By Day 21, the enrichment of aging-related pathways became more pronounced, with a modest increase in correlation with the AD transcriptome. This intermediate stage was characterized by the emergence of both metabolic decline and compensatory responses in SIRT6-regulated processes. At Day 30 time point, the correlation peaked, reflecting convergence on a gene expression profile more similar to that of AD, alongside widespread disruption of RNA metabolism, protein homeostasis, and neuronal structural pathways. Notably, SIRT6 re-expression at Day 30 Recovery led to a partial reversal of these transcriptomic changes, underscoring the potential of our model to identify pathways responsive to early intervention and potentially therapeutically targeted to prevent progression toward a pathological aging state. Moreover, here we demonstrated the flexibility of the shSIRT6 system, which initially captures an aging-like transcriptional state but, upon prolonged SIRT6 suppression, shifts toward a more pathological and AD-like profile, enabling the identification of pathways that transition from positive correlation in aging to anti-correlation in AD and providing a versatile framework to dissect molecular events underlying the progression from adaptive aging-associated responses to maladaptive disease-linked changes.

Together, our findings position the inducible SIRT6 silencing system as a powerful model for studying the temporal dynamics of aging. By mimicking the gradual decline of SIRT6, this system captures progressive molecular alterations that closely reflect those observed in aged and AD brains but also allows us to distinguish aging from AD signatures. Importantly, by enabling the distinction between reversible and irreversible changes, the recovery phase in our model not only deepens our understanding of how SIRT6 governs cellular aging but also provides a valuable framework for identifying candidate processes for chemical interventions and windows of opportunity to prevent the progression toward pathological aging.

## Acknowledgements and funding

The study was funded by the European Research Council (ERC) under the European Union’s Horizon 2020 research and innovation program (grant agreement No 849029), the David and Inez Myers foundation, the Israeli Ministry of Science and Technology (MOST), the Israel Science Foundation (ISF) 422/23, the High-tech, Bio-tech and Negev fellowships of Kreitman School of Advanced Research of Ben-Gurion University, The Israel Science Foundation (No. 277/22 DT). RNA-seq data analysis was partially supported by the Russian Science Foundation [25-71-20017 to E.K.]

## Methods

### Generation of shSIRT6 cells and SIRT6KO cells

SH-SY5Y cells were infected with the lentivirus Tet on/off shSIRT6 system, shRNAs targeting SIRT6, and a scrambled shRNA as a control. Cells were selected by 2 μg/mL puromycin for a week. To induce a gradual decline in SIRT6, cells were treated with DMEM supplemented with increasing doses of doxycycline (0.2 μg/mL, 0.5 μg/mL, 1 μg/mL) over a period of 30 days. During the recovery phase, doxycycline was eliminated from the cell media.

SH-SY5Y SIRT6KO or WT control cells were generated according to the protocol described in Kaluski et al., 2017^18^.

### Cell culture

All cells were grown in DMEM (catalog number: 41965039, Thermo-Fisher Gibco®, MA), supplemented with 1% L-glutamine (catalog number: 25030024, Thermo-Fisher Gibco®, MA), 1% Penicillin/Streptomycin antibiotics mix (catalog number: 15140122, Thermo-Fisher Gibco®, MA) and 10% FBS (catalog number: 12657-029, Thermo-Fisher Gibco®, MA). Incubation was 37°C, 5% CO2.

shSIRT6 cells media was supplemented with increased dosage of doxycycline (catalog number: 24390-14-5, Sigma-Aldrich) and with 1 μg/mL puromycin (catalog code: ant-pr-1, InvivoGen).

### RNA sequencing

#### RNA preparation

Cells were collected and total RNA was extracted using the NORGEN Total RNA purification kit (catalog number: 17250 according to the manufacturer’s manual.

#### RNA sequencing by Azenta

Total RNA samples were quantified using Qubit 4.0 Fluorometer (Life Technologies, Carlsbad, CA, USA) and RNA integrity was checked with 4200 TapeStation (Agilent Technologies, Palo Alto, CA, USA). Samples were treated with TURBO DNase (Thermo Fisher Scientific, Waltham, MA, USA) to remove DNA contaminants. RNA sequencing libraries were constructed based on ploy-A selection using Illumina value package, which was conducted following the manufacturer’s protocol.

The sequencing libraries were multiplexed and clustered on the flowcell. After clustering, the flowcell was loaded on the Illumina NovaSeq instrument according to manufacturer’s instructions. The samples were sequenced using 2×150bp, ∼350M PE reads (∼105GB), single index, value package.

### RNA-sequencing data analysis

#### RNA-seq data processing

RNA-seq read processing and alignment were conducted using nfcore/rnaseq (v3.7) pipeline^61^. Briefly, the quality of raw fastq files were evaluated with FastQC^62^, followed by read filtering using Trim Galore! (v0.6.7)^63^ and adapter removal with Cutadapt (v3.4)^64^. The strandedness of the samples was inferred via Salmon (v1.5.2)^65^ using *--libType A* setting. Processed reads were aligned to GRCh38 reference genome using STAR (v2.6.1d) tool^66^ and the transcript expression levels were estimated with Salmon using sorted and indexed BAM files generated by STAR as an input. These transcript-level expression estimates were then summarized at the gene level to produce a gene count matrix. Gene expression was normalized using DESeq2’s Median of Ratios method^67^. Principal component analysis was conducted using the *prcomp* function in R.

#### Differential expression analysis

To extract differentially expressed genes in shSIRT6 samples across days of doxycycline treatment (day 0, 5, 10, 21, 30), we applied likelihood ratio test (LRT) from DESeq2 with FDR p-value cut-off of 0.01. The resulting list of DEGs was further filtered to exclude genes that were significantly doxycycline-inducible in shCTRL replicates between day 0 and day 15, identified using the Wald test with an FDR p-value < 0.05. Differential expression analysis for pairwise comparisons between experimental days was performed using DESeq2 with adaptive shrinkage (*type = ‘ashr’*). Genes were considered significant if they met the criteria of FDR p-value < 0.05 and |log_2_(Fold Change)| > log_2_(1.5).

To perform differential analysis for day 30-specific comparisons, shSIRT6, shCTRL and shSIRT6-DMSO samples from two sequencing batches were combined into a single dataset. Low expressed genes were filtered out according to the following rule: more than 5 counts in at least 40% of samples, and zeros allowed in no more than 40% of samples. Differential expression analysis for day 30-specific comparisons was performed using the Wald test as described above, while controlling for batch effects by including batch as a covariate in the model: *∼ Batch + Experimental_group.* Differential expression analysis results were visualized as volcano plots using *EnhancedVolcano* (v1.22.0) R package^68^ and as an alluvial plot via *ggalluvia*l package (v0.12.5) To analyze genes that are specific to day 30, but have not fully returned to day 0 levels, an UpSet plot was generated to compare gene sets overlapping between stable genes from the comparisons of days 5, 10, and 21 with day 30. The plot was created using the ComplexUpset R package (v1.3.3).

### Analysis of age-related pathways

Lists of genes associated with various aging-related pathways (Ribosome biogenesis, RNA splicing, Glucose metabolic process, Protein folding, Oxidative phosphorylation, Cellular senescence, Cellular response to DNA damage) were obtained from the Gene Ontology database using *AnnotationDbi* (v1.66.0)^69^ and *org.Hs.eg.db* (v3.19.1)^70^ R packages. Gene list for “Oxidative phosphorylation” category was also supplemented with OXPHOS-related genes deposited in MitoCarta^69^ database. For the “Epigenetic regulation of gene expression” category, we combined “GOBP_EPIGENETIC_REGULATION_OF_GENE_EXPRESSION” and “REACTOME_EPIGENETIC_REGULATION_OF_GENE_EXPRESSION” (both v2023) gene sets, derived from mSigDB^71^. Genes that induce or inhibit senescence were retrieved from the CellAge database^72^, and only those associated with non-cancer cellular senescence were retained for analysis.

To visualize longitudinal trends of DE genes associated with each aging-related pathway, normalized counts were rlog-transformed, and the mean gene expression per day was calculated. Data were then standardized using z-transformation to allow comparison of expression trends. One-way ANOVA test was applied to assess the significance of gene expression changes across days of experiment for each pathway of interest.

### Gene expression clustering and clusters enrichment analysis

Clustering of the expression profiles of DE genes was carried out using the degPattern function from the *DEGreport* R package (version 1.40.1), with the following parameters: “minc = 15”, “cutoff = 0.7”, and “consensusCluster = FALSE”. Gene Ontology and KEGG enrichment analyses were performed via *clusterProfiler* (v4.12.6) package^73^. Reactome pathway analysis was conducted using the *Reactome PA* (v1.48.0) package^74^. For all the enrichment analyses, gene symbols were converted to ENTREZ IDs using *AnnotationDbi* (v1.66.0) and *org.Hs.eg.db* (v3.19.1) packages using all expressed genes as a background. Redundant GO categories were filtered out using the *simplify* function with the Wang method for semantic similarity implemented in *clusterProfiler*. Terms or pathways satisfying FDR p-value < 0.05 threshold (Fisher’s test) were retained as significantly altered.

### Comparative analysis with transcriptome of the human brain

#### GSEA NES Correlation Between Aged Brain Regions and Day21

Transcriptional changes associated with aging were investigated using GTEx Analysis v10 RNA-seq data and the microarray data GSE2521928, stratified by brain region. DEGs between old and young samples were identified using the DESeq2 (version 1.44.0) for RNA-seq data and limma package (version 3.60.6) for microarray data, applying a false discovery rate (FDR) threshold of < 0.05. Age-associated DEGs were then compared with those identified between Day21 and Day0 in our dataset. Pathway enrichment analysis was performed using MSigDB gene sets (C2, C3, C5) via the fgsea package (v1.30.0), with pathway interest defined as those with FDR < 0.2. Overlapping enriched gene sets between Day21 and each brain region were used to generate scatter plots for visualization. The correlation between old and Day21 was assessed by linear regression analysis with the model: Old ∼ Day21.

### Comparative analysis with human transcriptome portraits of AD

#### Intersection Plot (Modified UpSet Plot)

Human middle-aged and Alzheimer’s disease (AD) brain samples were obtained from the GSE153875 dataset^29^ and integrated with SH-SY5Y cell line samples based on gene symbol matching. Data normalization was carried out using genes that exhibited stable count expression (coefficient of variation < 0.1) across all samples. Differentially expressed genes (DEGs) between AD and middle-aged samples were identified using DESeq2 (version 1.44.0) with a stringent significance threshold (adjusted p-value ≤ 0.01). The resulting DEGs were analyzed for intersections across experimental conditions (AD vs. middle age; Day10 vs. Day0; Day21 vs. Day0; Day30 vs. Day0; Day30 recovery vs. Day0). Visualization was performed using R scripts (base version 4.4.0 and ggplot2 v)

### GSEA NES Correlation Between Alzheimer’s Disease and time points

Differentially expressed genes (DEGs) for each time point were identified from the following comparisons: Day10 vs. Day0, Day21 vs. Day0, Day30 vs. Day0, and Day30 recovery vs. Day0. For Alzheimer’s disease (AD)-specific analysis, DEGs were obtained from AD vs. middle-aged comparison. Gene set enrichment analysis (GSEA) was conducted using the Molecular Signatures Database (MSigDB; C2, C5, and C5 collections) via fgsea package. Enriched gene sets were filtered at a significance threshold of Normalized Enrichment Score (NES) FDR < 0.05 for downstream analysis. To assess overlap between AD and temporal profiles, scatter plots were generated from shared gene sets between AD and each time point.

To evaluate the association between AD and the time points, we employed a combination of Pearson correlation, permutation testing, and bootstrapping. The Pearson correlation coefficient was calculated to quantify the linear relationship between AD and time points. Statistical significance was assessed using a permutation test, and a 95% confidence interval (CI) for the correlation coefficient was estimated via bootstrapping.

To summarize the association with single metric, we applied Fisher’s z transformation to Pearson’s r in to z = atanh(r), then standardized it by its standard error, 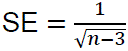 gives,

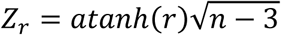

This Zr is a standardized effect size: it keeps the sign of the correlation, and its magnitude reflects both the strength of the association and its precision.

#### Circos plot

To identify differential enrichment of functional pathways, GSEA-derived gene sets from AD and aged samples were compared. Pathways demonstrating shifts in normalized enrichment scores were visualized using the circlize package (version 0.4.16) in R.

#### Bubble Plot (GO Enrichment Analysis of Overlapping DEGs)

To identify biological pathways shared between AD and experimental time points, DEGs were stratified into upregulated (log₂FC > 0) and downregulated (log2FC < 0 categories for each comparison (AD, Day10, Day21, Day30, and Day30 recovery). For each time point, condition-specific gene sets were defined as intersections with AD-associated DEGs: (1) AD ∩ Day10, (2) AD ∩ Day21, (3) AD ∩ Day30, and (4) AD ∩ Day30 recovery. Gene Ontology (GO) enrichment analysis was performed using the same methods in clusterProfiler, with significance defined at FDR < 0.05. Dot plots were generated using ggplot2 (version 3.5.2).

### Protein extraction

#### Chromatin-bound protein acid extraction

The chromatin-bound proteins were extracted from cell culture dry pellets according to the following protocol: Cells pellets were resuspended in cytoplasmic protein lysis buffer (described below), in a volume equivalent to 3 times the volume of the pellet and homogenized by thorough pipetting. Once homogenized, samples were kept on ice for 20 minutes, then centrifuged for 10 minutes, 21100g, 4°C. The supernatant, which contains the non-chromatin-bound fraction of proteins, was transferred into new tubes. The pellets (which contain the chromatin and cell leftovers) were washed twice with the same volume of cytoplasmic protein lysis buffer, incubated for 5 minutes on ice and centrifuged for 5 minutes, 21100g, 4°C. then, Add 0.2N HCl solution to the dry pellet (equivalent of 1/8-1/10 of the original cytoplasmic protein lysis buffer volume) and pipette thoroughly. Samples were incubated on ice for 20 minutes with occasional vortexes, then centrifuged for 10 minutes, 21100g, 4°C. Supernatants (contain the chromatin-bound proteins) were transferred to new tubes and neutralized by adding 1M Tris pH8 (the same volumes as the 0.2N HCl), then vortexed. Protein concentrations were determined using Bradford assays.

cytoplasmic protein lysis buffer: 10 mM HEPES pH7.4, 10mM KCl, 0.05% NP-40, phosphatase inhibitor cocktail X1 (APExBIO K1013), 0.2mM PMSF.

#### Nuclear and cytoplasmic protein extraction

The Nuclear and cytoplasmic proteins were extracted from cell culture dry pellets according to the following protocol: Cells pellets were resuspended in CHB buffer (described below), in a volume equivalent to 3 times the volume of the pellet and homogenized by thorough pipetting. Once homogenized, samples were kept on ice for 15 minutes, then 1% NP-40 was added to the samples. Samples were then centrifuged for 10 minutes, 700g, 4°C. The supernatant, which contains the cytoplasmic fraction of proteins, was transferred into new tubes.

The pellets (which contain the nuclear proteins) were resuspended in full RIPA lysis buffer (described below), incubated 30 minutes on ice with occasional vortexes and centrifuged for 15 minutes, 21100g, 4°C. Supernatants were transferred to a new tube.

Protein concentrations were determined using Bradford assays.

CHB buffer: 20mM Tris pH7.4, 10mM NaCl, 3mM MgCl_2_, phosphatase inhibitor cocktail X1 (APExBIO K1013), 0.2mM PMSF.

Full RIPA buffer: 5mM Tris pH7.4, 15mM NaCl, 1mM EDTA, 1% NP-40, 0.1% SDS, 0.5% sodium deoxycholate, phosphatase inhibitor cocktail X1 (APExBIO K1013), 0.2mM PMSF.

### Western blots

For each Western blot analysis, 10-40 µg protein samples were loaded onto 8–15% Tris-glycine polyacrylamide gel gels made in house. Proteins were separated for 1 h at 120 V and then blotted to nitrocellulose membranes at 100 V for 90 min. The blots were blocked with 5% skim milk in TBST (15 mM Tris-HCl, pH 7.5, 200 mM NaCl, and 0.1% Tween 20) for 1 h at room temperature. Membranes were incubated overnight with primary antibodies, diluted per manufacturer recommendation and developed using a chemiluminescence reagent (WesternBright Quantum HRP substrate, K-12042-C20).

### Immunofluorescence

Cells were rinsed with phosphate buffer saline (PBS) and fixed with 4% paraformaldehyde for 10 min at room temperature, followed by two additional washes. Cells were permeabilized (0.1% tri-sodium citrate and 0.1% Triton X-100 in Distilled water, pH 6) for 5 min and rinsed again. After 40 min of blocking (0.5% bovine serum albumin [BSA], 5% goat serum, 0.1% Tween-20 in PBS), cells were incubated with primary antibody diluted in blocking buffer overnight at 4°C. The next day, cells were washed three times with wash buffer (0.25% BSA, 0.1% Tween-20 in PBS), incubated for 1h with the secondary antibody (diluted in blocking buffer 1:200) at room temperature and rinsed three more times. Cells were then Hoechst-stained for three minutes at room temperature and rinsed with PBS twice before imaging.

### Image analysis and quantification

Nuclear mean intensity was measured and analyzed using ImageJ (https://imagej.net/ij/download.html).

For quantification of nuclear size and foci count, we used CellProfiler 4.2.6. The cell nucleus was first segmented based on shape and intensity using the Hoechst channel and used as primary object reference. For the foci count, a module was added to identify objects within the nucleus according to typical diameter in pixels units. This enabled the quantification of the desired parameter (total nuclear area, intensity and foci count per nucleus) while eliminating objects outside the nucleus. To verify segmentation, we asked CellProfiler to overlap each channel with the respective objects and masks, generating images for quality control. Only cells with accurate nucleus and nucleolus segmentation (validated by overlap images) were included in the statistical analysis, with inaccurately segmented cells excluded. A detailed spreadsheet with curated data was generated for each experiment.

### Evaluation of Lamin B abnormalities

Evaluation of Lamin B abnormalities accumulation during SIRT6 silencing was done by setting 4 parameters per cell: circularity, broken envelope, mislocalization and foci of Lamin B staining, giving a score of one or zero for each parameter.

### Lentiviral Transduction

To enable live-cell visualization of nuclear and cytoplasmic compartments, SH-SY5Y cells were transduced with a lentiviral vector encoding a GFP-tagged nuclear export signal (GFP:NES) and an RFP-tagged nuclear localization signal (RFP:NLS) (Addgene Plasmid #71396). Lentiviral particles were generated in HEK293T cells using a third-generation packaging system. Viral supernatants were collected 48 hours post-transfection, centrifuged, filtered through 0.45 μm PVDF membranes, and directly applied to SH-SY5Y cultures. Transduction efficiency was evaluated 72 hours post-infection by fluorescence microscopy.

#### Fluorescence-Activated Cell Sorting (FACS)

Transduced SH-SY5Y cells were resuspended in sterile FACS buffer (PBS supplemented with 2% FBS and 1 mM ethylenediaminetetraacetic acid (EDTA)) and subjected to fluorescence-activated cell sorting. Cells co-expressing GFP and RFP were isolated using a BD FACSAria™ III cell sorter (BD Biosciences) and subsequently expanded for downstream applications.

### Statistical analysis

Between-group comparisons that used t-test, ANOVA (one-way or two-way, with post hoc Dunnet or Tukey tests), and Wilcoxson rank sum test were conducted at the significance level of 0.05 and performed using GraphPad Prism 7. Population proportion analysis was performed using the chi-square test to assess differences in categorical variables between groups.

---

**Supplementary Figure 1.**
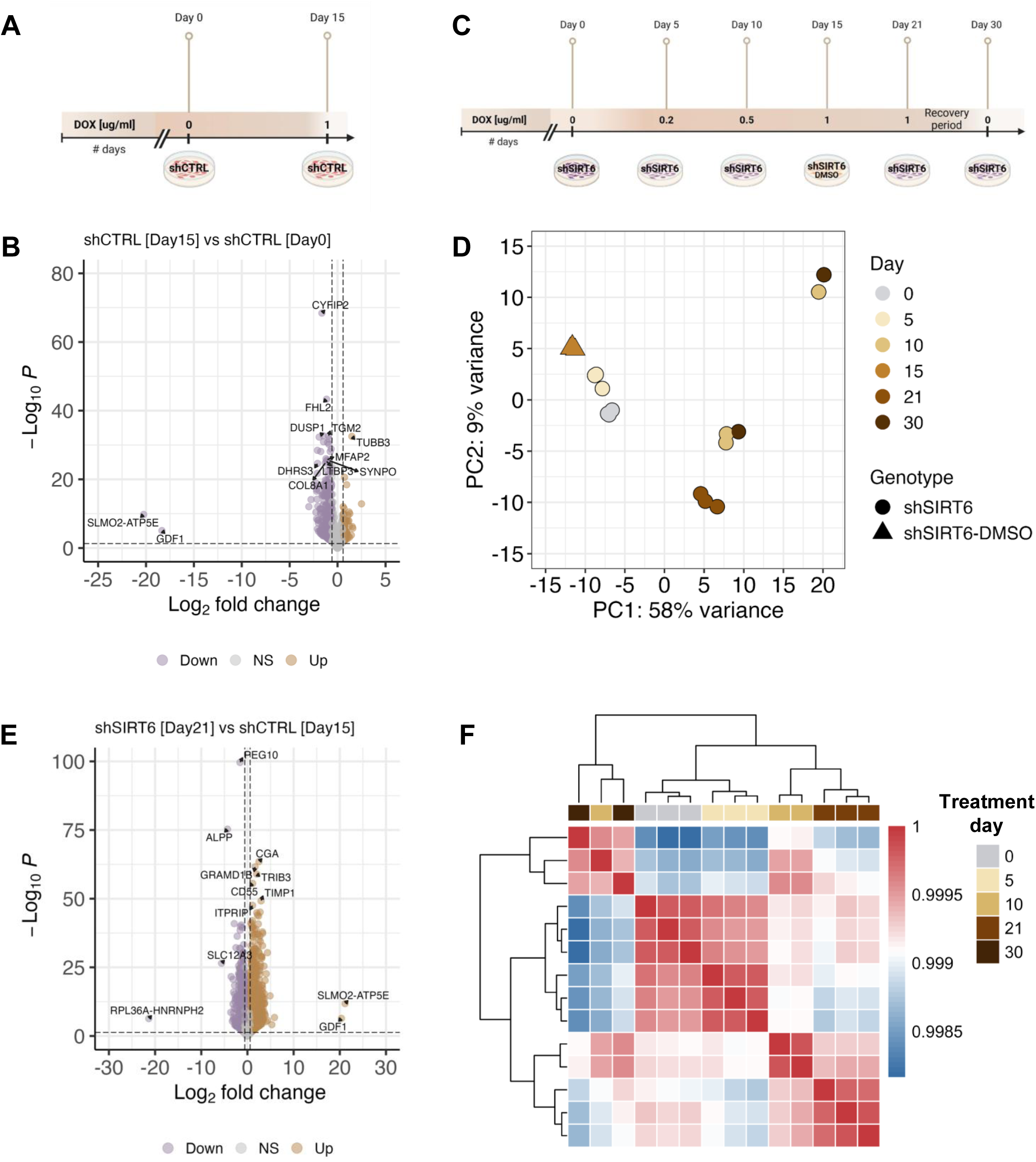
(**A**) Schematic representation of untreated and doxycycline (Dox)–treated scrambled shRNA (shCTRL) cells. **(B)** Differentially expressed genes between untreated and Dox-treated shCTRL cells, defined as off-target/Dox effect genes. **(C)** Schematic representation of the experimental design using the shSIRT6 cell line, with shSIRT6 cells treated with DMSO as a mock control. **(D)** Principal component analysis illustrating the similarity among replicates. **(E)** Differentially expressed genes between shSIRT6 cells treated with Dox for 21 days and shCTRL cells treated with Dox for 15 days, reflecting changes associated with SIRT6 silencing. **(F)** Heatmap showing sample clustering based on pairwise Pearson correlation coefficients, with red indicating high correlation and blue indicating low correlation.

**Supplementary Figure 2.**
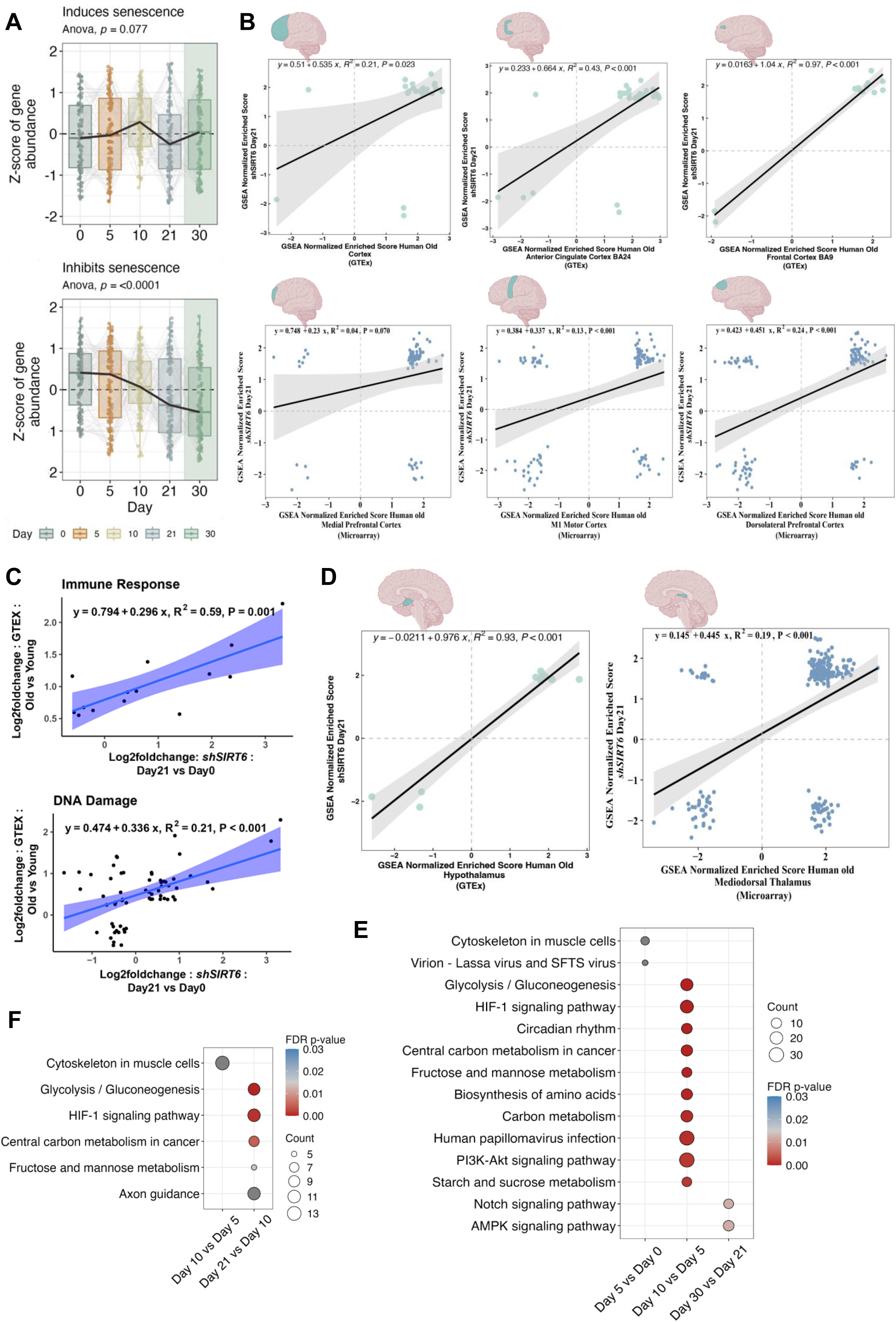
(**A**) Expression trends of senescence-related gene inducers and inhibitors during SIRT6 silencing (Days 0, 5, 10, and 21) and after recovery (Day 30, green background). **(B)** Correlation plot of gene sets enriched in aging human cortical brain regions based on GTEx (upper) and microarray (lower) datasets and those enriched in shSIRT6 Day 21 cells. **(C)** Gene-level scatter plot of selected pathways from the GTEx hippocampus and Day 21 comparison. **(D)** Correlation plot of gene sets enriched in the human hypothalamus (left) and thalamus (right) aging brain and those enriched in shSIRT6 Day 21 cells. **(E)** Pathway enrichment analysis of upregulated genes between consecutive time points. **(F)** Pathway enrichment analysis of downregulated genes between consecutive time points. Colors indicate FDR-adjusted p values (hypergeometric test, p < 0.05), and circle size is proportional to gene count.

**Supplementary Figure 3.**
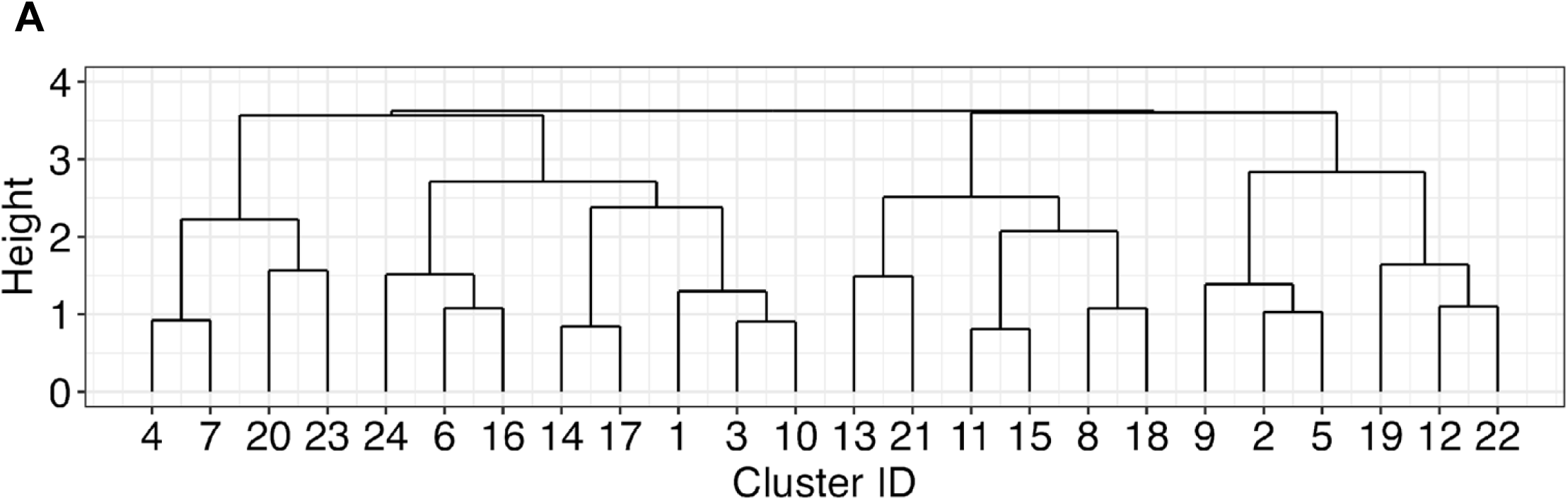
(**A**) Representation of the 24 DE genes clusters, with annotations indicating gene cluster similarity obtained using hierarchical clustering.

**Supplementary Figure 4.**
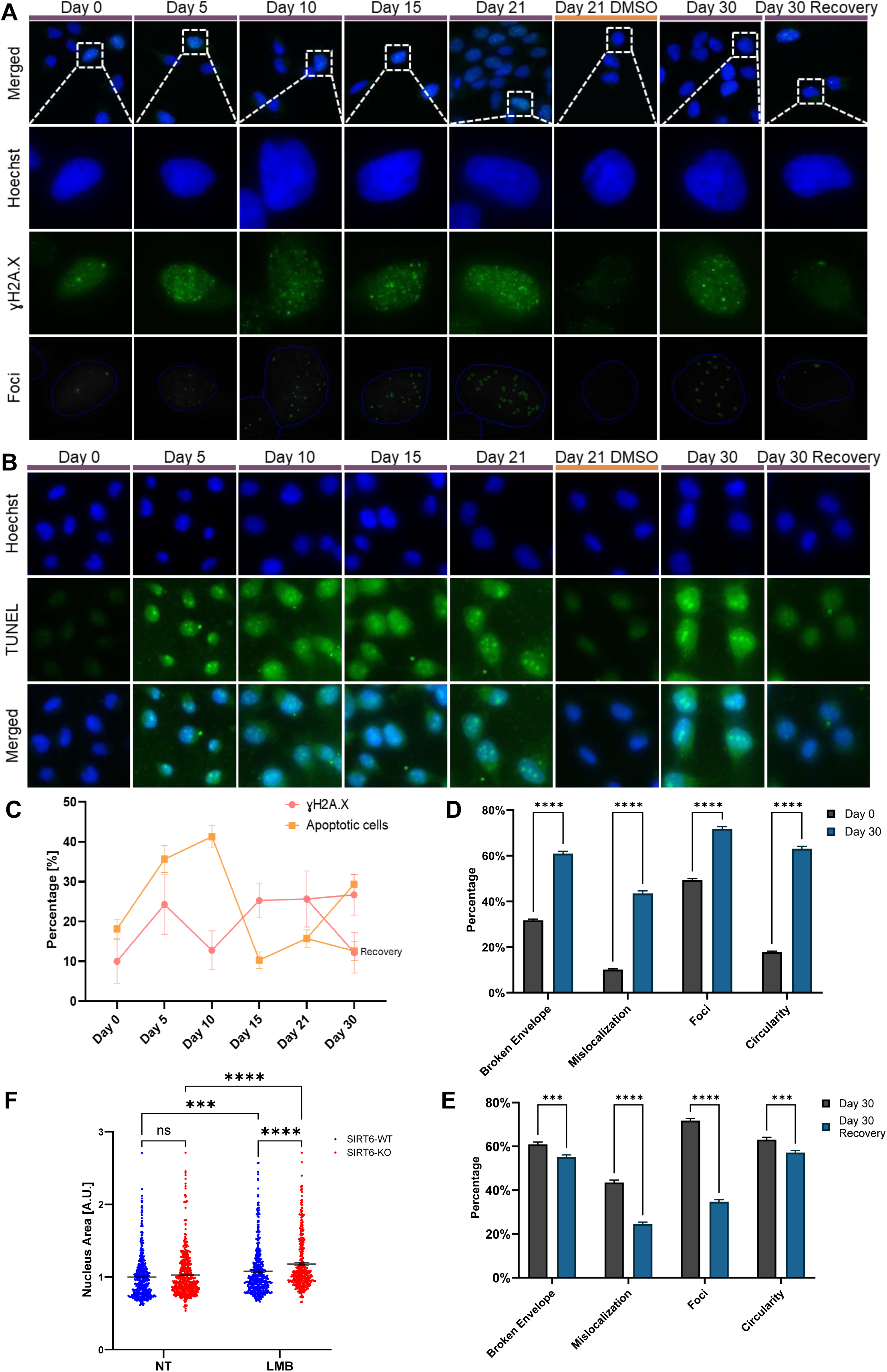
(**A**) Representative immunofluorescence images showing the double-strand break marker γH2A.X in cells subjected to SIRT6 modulation. **(B)** Representative immunofluorescence images of TUNEL staining used to assess cell viability in cells subjected to SIRT6 modulation. **(C)** Line plot showing mean percentages over time of γH2A.X-positive cells (pink) and apoptotic cells (orange). **(D,E)** Relative fold change of four parameters used to assess nuclear integrity comparing Day 0 and Day 30 (D) and between the two conditions at Day 30 (E). **(F)** Nuclear area measurements in SIRT6 wild-type (WT) and knockout (KO) cells. Data are presented as mean ± SEM (ANOVA; NS, not significant; *P < 0.05, **P < 0.01, ***P < 0.001, ****P < 0.0001).

**Supplementary Figure 5.**
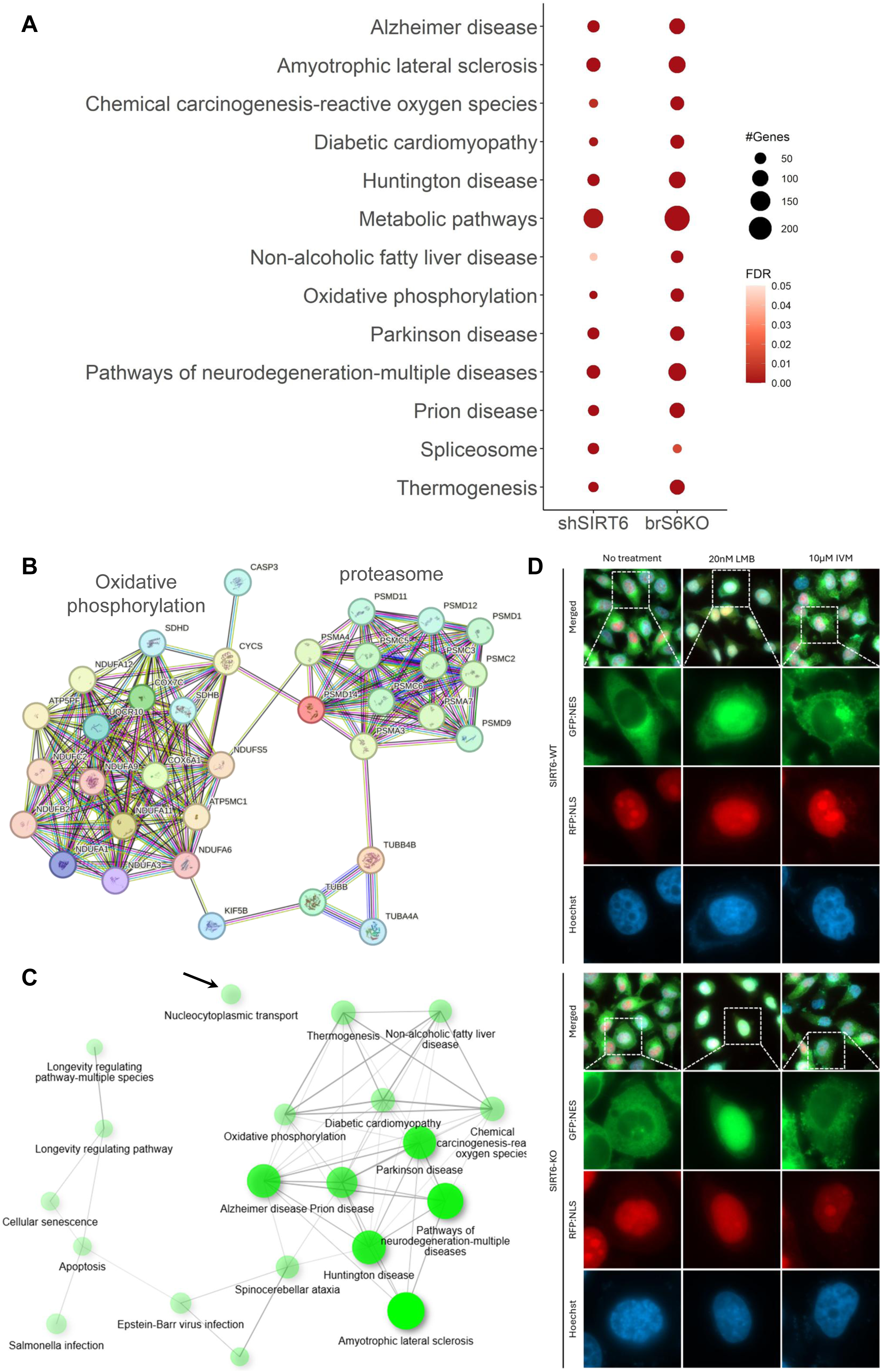
(**A**) Overlap of enriched pathways between brS6KO and shSIRT6 cells. Colors indicate FDR-adjusted p values, and circle size is proportional to gene count. **(B)** Molecular function network of 34 genes shared across five neurodegenerative diseases. **(C)** Gene network of 112 neurodegenerative disease–related genes, with the black arrow highlighting nucleocytoplasmic transport (NCT) as a distinct network category. (D) Representative immunofluorescence images of the GFP:NES–i–RFP:NLS reporter in untreated SIRT6 wild-type (WT) and knockout (KO) cells and after 24 h treatment with LMB or IVM.

**Supplementary Figure 6.**
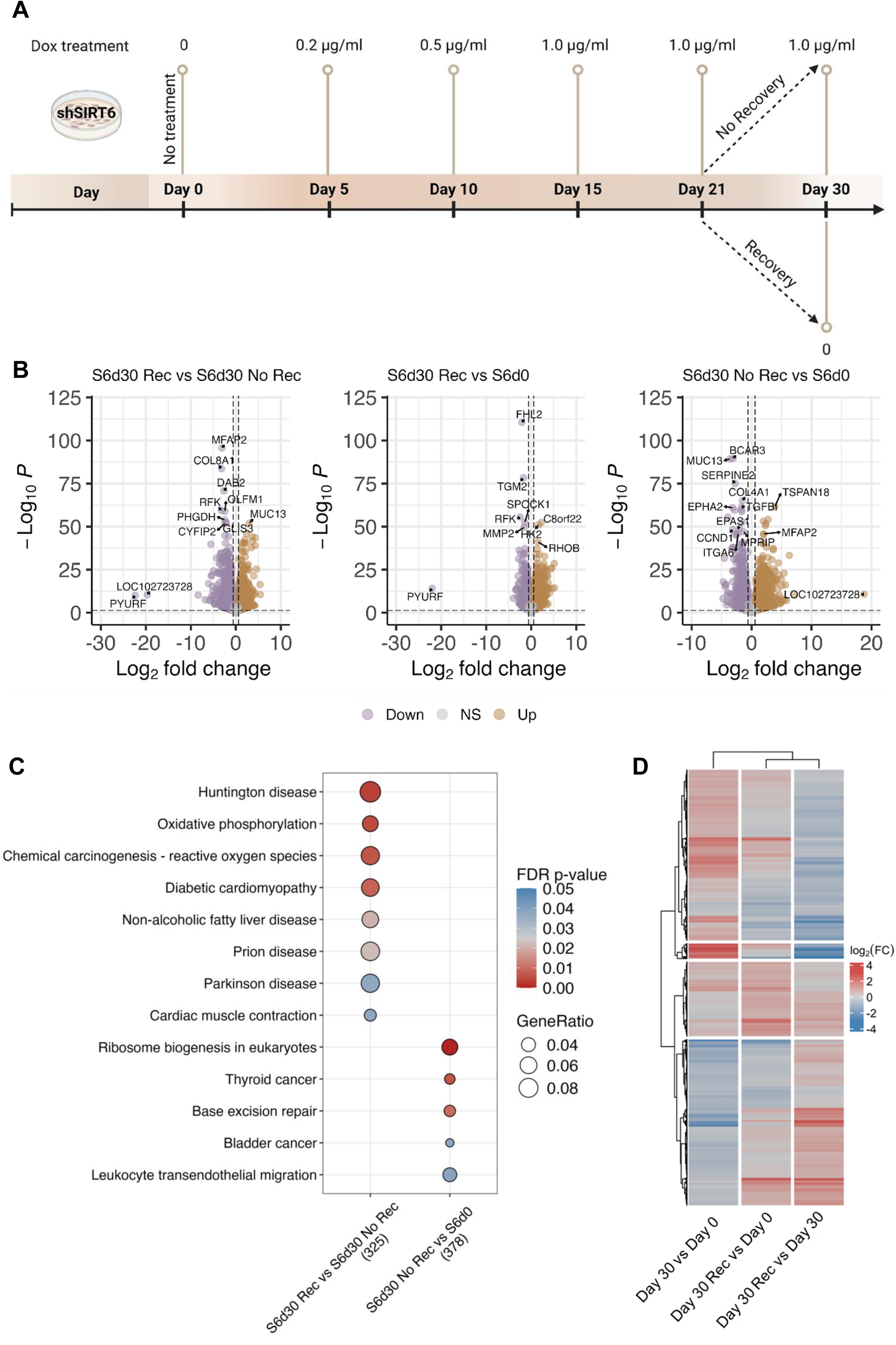
(**A**) Schematic overview of the shSIRT6 aging-like model, highlighting the two experimental conditions at the Day 30 time point. **(B)** Volcano plots of differentially expressed genes from three comparisons: Day 0 vs. Day 30, Day 0 vs. Day 30 recovery, and Day 30 vs. Day 30 recovery. **(C)** KEGG pathway enrichment analysis performed separately on the non-overlapping DEGs from the Day 0 vs. Day 30 and Day 30 vs. Day 30 recovery comparisons. **(D)** Heatmap of 327 shared genes, showing their separation into four distinct subgroups based on expression trends.

**Supplementary Figure 7.**
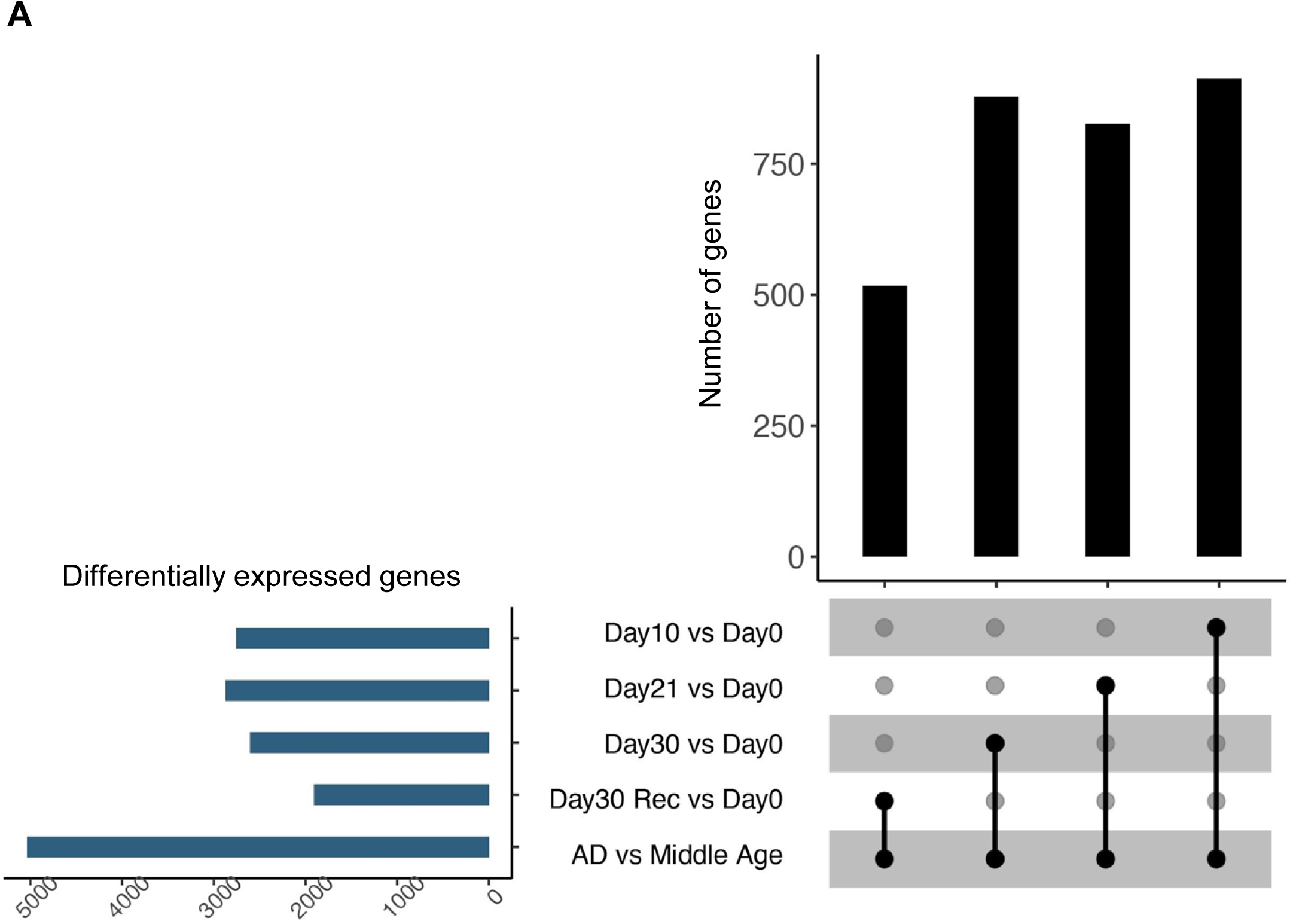
(**A**) UpSet plot presenting overlap of the genes changed in human AD patients and in shSIRT6 cells at different time points.

